# Ribosome stalling during selenoprotein translation exposes a ferroptosis vulnerability in cancer

**DOI:** 10.1101/2022.04.11.487892

**Authors:** Zhipeng Li, Lucas Ferguson, Kirandeep K. Deol, Melissa A. Roberts, Leslie Magtanong, Michael C. Bassik, Scott J. Dixon, Nicholas T. Ingolia, James A. Olzmann

## Abstract

Ferroptosis is a regulated, iron-dependent form of necrosis that is triggered by the accumulation of oxidatively damaged phospholipids^1–3^. Glutathione peroxidase 4 (GPX4) prevents ferroptosis by converting phospholipid hydroperoxides into non-toxic lipid alcohols^4, 5^. Ferroptosis has been implicated in the pathology of several degenerative conditions and inhibiting GPX4 activity has emerged as a therapeutic strategy to induce cancer cell death^1, 2^. However, many cancer cell lines are resistant to GPX4 inhibition^6^, and the mechanisms that regulate GPX4 activity and ferroptosis resistance remain incompletely understood. Here, employing a synthetic lethal CRISPR-Cas9 screen in a triple negative breast cancer (TNBC) cell line, we identify LRP8 (also known as ApoER2) as a ferroptosis resistance factor. LRP8 is upregulated in cancer, and we find that it promotes ferroptosis resistance in cancer cells in both 2-dimensional (2-D) cell culture and 3-dimensional (3-D) spheroid models. Mechanistically, loss of LRP8 decreases cellular selenium levels, resulting in the reduced expression of a subset of selenoproteins, including GPX4. Remarkably, the reduction in GPX4 is not due to the classic hierarchical selenoprotein regulatory program^7, 8^. Instead, our findings demonstrate that the translation of GPX4 is severely impaired in the selenium-deficient LRP8 knockout (KO) cells due to extensive ribosome stalling at the inefficiently decoded GPX4 selenocysteine (SEC) UGA codon, which results in ribosome collisions and early translation termination. Thus, our findings reveal ribosome stalling and collisions during GPX4 translation as targetable ferroptosis vulnerabilities in cancer cells.

## MAIN

Two primary cellular mechanisms that prevent ferroptosis are the conversion of phospholipid hydroperoxides into non-toxic lipid alcohols by GPX4^4^ and the generation of radical trapping antioxidants to block the propagation of lipid peroxidation by enzymes such as ferroptosis suppressor protein 1 (FSP1)^9, 10^, dihydroorotate dehydrogenase (DHODH)^11^, and GTP cyclohydrolase-1 (GCH1)^12, 13^. Targeting the pathways that mediate ferroptosis resistance has emerged as a promising cancer therapeutic strategy, but how known protective pathways are regulated and whether additional mechanisms of ferroptosis resistance exist remain outstanding questions.

### LRP8 is a candidate ferroptosis regulator that is upregulated in cancer

High *FSP1* expression is strongly correlated with resistance to GPX4 inhibitors^9, 10^, consistent with the role of FSP1 in preventing ferroptosis. To identify additional ferroptosis resistance mechanisms, we searched for cell lines displaying FSP1-independent resistance. Analysis of cancer cell line sensitivities to ferroptosis inducers reported in the Cancer Therapeutics Response Portal (CTRP)^14, 15^ identified a subset of cancer cell lines, including the MDA-MB-453 TNBC cell line, that are resistant to GPX4 inhibitors (RSL3, ML162, and ML210) despite low amounts of FSP1 mRNA (**Fig. 1a and Extended Data Fig. 1a,b**). Consistent with these data and in contrast to U-2 OS cells, MDA-MB-453 cells lack detectable FSP1 protein (**Fig. 1b**), and treatment with the FSP1 inhibitor iFSP1 (**Fig. 1c**) or expression of FSP1 sgRNAs (**Extended Data Fig. 1c**) have no effect on RSL3-induced cell death. These findings indicate that MDA-MB-435 cells employ FSP1-independent mechanisms to prevent cell death triggered by RSL3.

**Fig. 1:**
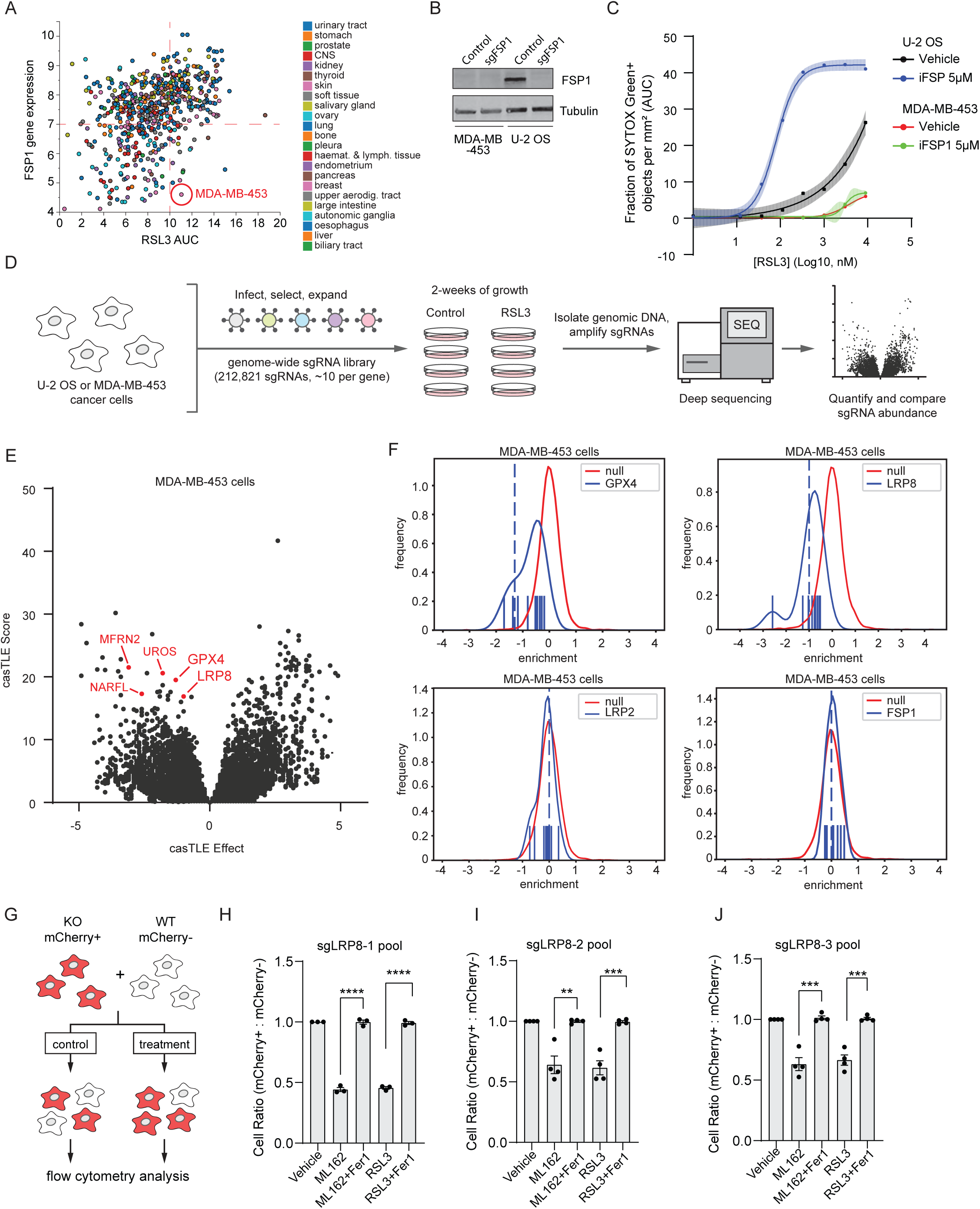
Genome-wide CRISPR screen identifies LRP8 as a genetic modifier of ferroptosis resistance in triple-negative breast cancer cells. **a**, Summary of FSP1 gene expression and RSL3 sensitivity in various cancer cells, data were mined from the CTRP database. AUC is the area under the curve from RSL3 dose response analyses of cell viability. **b**, Western blot of lysates from MDA-MB-453 cells and U2-OS cells indicates FSP1 protein levels and KO efficiency. **c**, Dose response of RSL3 induced cell death in U-2 OS and MDA-MB-453 cells treated with vehicle or 5µM iFSP1. Shading indicates 95% confidence intervals for the fitted curved and each data point is the average of three replicates. AUC, Area Under the Curve. **d**, Schematics of CRISPR-Cas9 screen strategy. **e**, Gene effects and gene scores calculated for individual genes analyzed in the genome-wide CRISPR screen in MDA-MB-453 cells. **f**, Histograms of selected individual gene results from casTLE analysis, including the average for the entire population (red line) and for the indicated gene sgRNAs (blue line). **g**, Schematic of cell competitive growth assay. **h-j**, Quantification of the ratio between mCherry+ : mCherry-cells after the indicated treatment. Data represent mean ± S.E.M. of three biological replicates, which were compared using a two-tailed, unpaired t-test.

To identify new mechanisms of ferroptosis resistance, we performed parallel genome-wide synthetic lethal CRISPR-Cas9 screens with RSL3 in FSP1-expressing U-2 OS cells and FSP1-deficient MDA-MB-453 cells (**Fig. 1d**). Consistent with our previous CRISPR-Cas9 screen with a sublibrary of sgRNAs^9^, FSP1 emerged as the top ferroptosis suppressor in U-2 OS cells (**Extended Data Fig. 1d,e and Supplementary Table 1**), indicating that FSP1 mediates a primary mechanism of ferroptosis resistance in response to GPX4 inhibition in these cells. As expected, *FSP1* was not identified as a ferroptosis regulator in MDA-MB-453 cells (**Fig. 1e,f and Supplementary Table 2**). However, several other regulators of ferroptosis resistance were observed, including *GPX4* and factors involved in iron regulation, such as enzymes that mediate heme synthesis and iron-sulfur cluster formation (e.g., *MFRN2*, *UROS*, *NARFL*) (**Fig. 1e,f and Supplementary Table 2**). Interestingly, sgRNAs targeting *LRP8*, which belongs to the lipoprotein receptor superfamily, were substantially disenriched in MDA-MB-453 cells treated with RSL3 (**Fig. 1e,f and Supplementary Table 2**). In addition, treatment of co-cultured wildtype (WT) cells (mCherry-) and cells expressing three different LRP8 sgRNAs (mCherry+) with the GPX4 inhibitors RSL3 or ML162 consistently showed a reduced mCherry+/mCherry-ratio in a ratiometric competitive growth assay (**Fig. 1g-j**). This growth disadvantage was reversed by culturing the cells in the presence of the ferroptosis inhibitor ferrostatin-1 (Fer1) (**Fig. 1h-j**), suggesting that the loss of LRP8 sensitizes cells to ferroptosis.

LRP8 also stood out as an interesting hit in our screen due to strong functional connections with cancer. Analysis of data in The Cancer Genome Atlas (TCGA) indicates that *LRP8* is upregulated in tumors from multiple primary sites, including breast tissue (**Fig. 2a and Extended Data Fig. 2a**), and patients with higher expression of *LRP8* have poorer survival rates (**Extended Data Fig 2b**). Furthermore, in data collected from over 800 cancer cell lines in the CTRP^14, 15^, LRP8 expression is positively correlated with resistance to the GPX4 inhibitors RSL3, ML162, and ML210 (**Fig. 2b**). These data suggest that LRP8 promotes cancer growth and mediates ferroptosis resistance across a broad range of cancer cells.

**Fig. 2:**
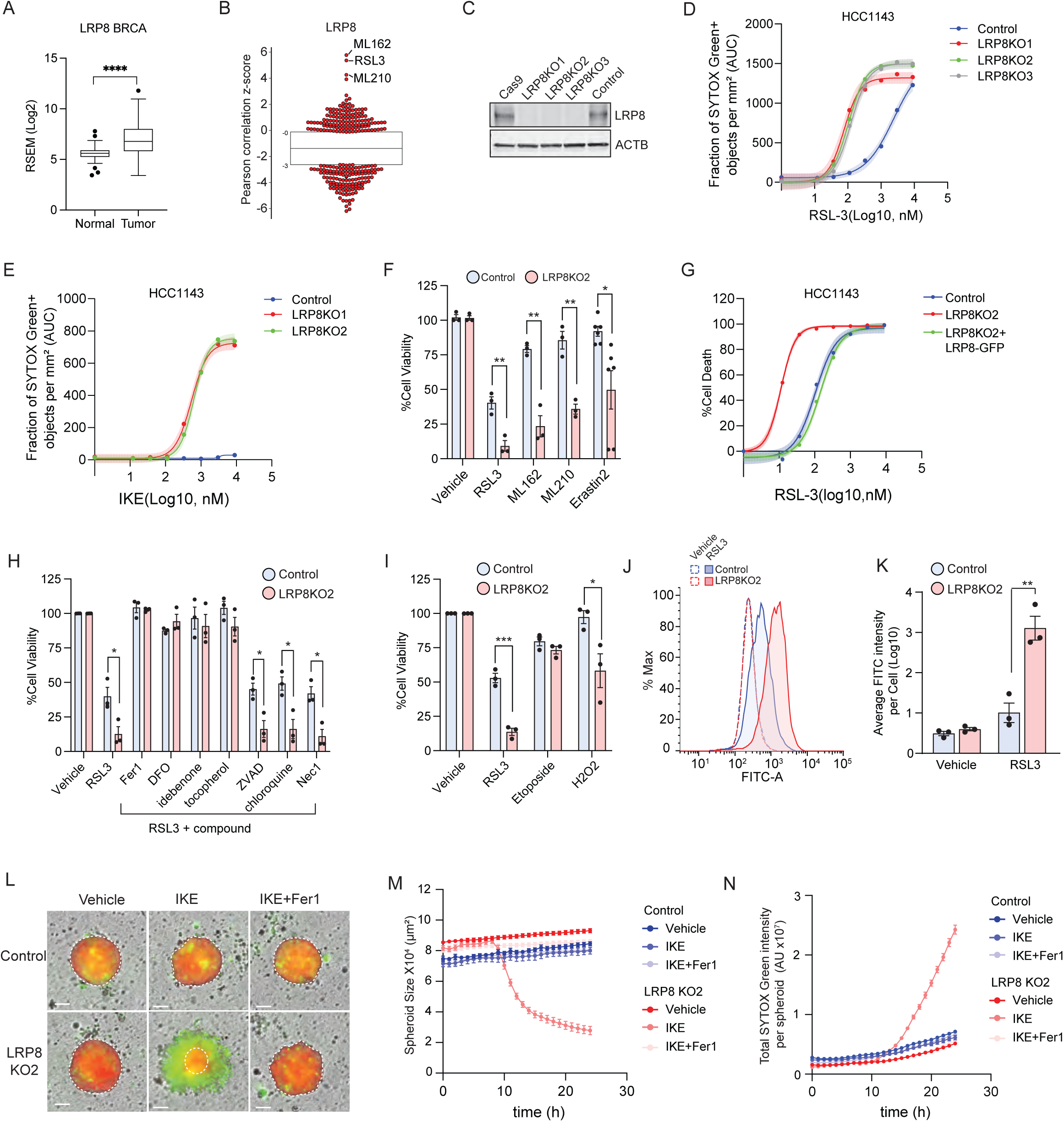
LRP8 promotes ferroptosis resistance in breast cancer cells. **a**, Relative expression of LRP8 in normal and breast tumor tissue. Data was mined from the TCGA-BRCA database. **b**, LRP8 expression is positively correlated with resistance to ferroptosis inducers in cancer cells. **c**, Western blot analysis of the LRP8 KO in HCC1143 cells with multiple sgRNAs. **d**, Dose response of RSL3-induced cell death, SYTOX Green (SG) positive (SG+) cells, in sgSAFE control (referred as “control” hereafter) and LRP8 single clonal KO cells. **e**, Dose response of IKE-induced cell death in control and LRP8KO cells. **f**, Cell viability of control and LRP8KO cells 24 hr after treatment with the indicated ferroptosis inducers by cell-titer glow assay (RSL3, 200nM; ML162, 200 nM; ML210, 400 nM; Erastin 2, 400 nM). **g**, Dose response of RSL3-induced cell death in control, LRP8KO, and LRP8KO expressing LRP8-GFP by cell titer glow assay. **h**. Cell viability of control and LRP8KO cells treated with RSL3 in the presence of inhibitors of different types of cell death for 24 hr (RSL3, 200 nM; Fer1, 2 µM; DFO, 100 µM; idebenone, 10 µM; tocopherol, 10 µM; ZVAD, 20 µM; chloroquine, 200 µM; Nec-1, 1 µM). **i**, Cell viability of control and LRP8KO cells following treatment with RSL3, etoposide, and H_2_O_2_ for 24 hr (RSL3, 200 nM; etoposide, 100 µM; H_2_O_2,_ 30 µM). **j**, Control and LRP8KO cells treated with 200nM RSL3 for 5 hrs were labeled with BODIPY 581/591 C11. Green fluorescence intensity was analyzed by flow cytometry. **k**, Quantification of FITC intensity from **j**. **l**, Spheroids from control and LRP8KO cells stably expressing mCherry were incubated with IKE or IKE+Fer1 and SG and visualized using live-cell imaging. **m**,**n**, Quantification of the size of each spheroid (**m**) and the total intensity of the SG signal (**n**) over 24 hr. In **d-f**, Shading indicates 95% confidence intervals for the fitted curves and each data point is the average of three biological replicates. All data represent mean ± S.E.M. of three biological replicates, except for Erastin2 in **f** (n=6), by two-tailed, unpaired t-test (**f, h, i, k**). For **m** and **n**, at least 8 independent spheroids were collected in three biological repeats.

### LRP8 promotes ferroptosis resistance in cancer cells

To experimentally evaluate the relationship between LRP8 and ferroptosis, we generated multiple clonal LRP8 KO cell lines in MDA-MB-453, HCC1143, and HCC1937 TNBC cell lines (**Fig. 2c and Extended Data Fig. 2c,d**). Quantification of cell viability using time-lapse microscopy and the Cell-Titer Glo assay revealed a substantial increase in the sensitivity of LRP8KO TNBC cells to RSL3 (**Fig. 2d and Extended Data Fig. 2e-h**). Similarly, depletion of LRP8 sensitized a panel of other cancer cell lines to RSL3, including the glioblastoma cell line U87-MG and the melanoma cell lines A375 and SK-MEL28 (**Extended Data Fig. 3a-d**), indicating a generalizable role as a ferroptosis resistance factor in cancer cells. Moreover, LRP8KO cells were sensitized to ferroptosis induced by depletion of glutathione with the system x_c_^-^ inhibitors Erastin2 and imidazole ketone erastin (IKE) as well as by two additional GPX4 inhibitors (ML162 and ML210) (**Fig. 2e,f**). The LRP8KO sensitivity to RSL3 and IKE was reversed by re-expression of LRP8-GFP (**Fig. 2g and Extended Data Fig. 2i-k**), indicating that the phenotype is due to the loss of LRP8. RSL3-induced cell death in LRP8KO cells was blocked by several known ferroptosis inhibitors, including radical trapping/scavenging antioxidants, such as Fer1, idebenone, and tocopherol, and the iron chelator deferoxamine (DFO) (**Fig. 2h and Extended Data Fig. 2l**), but not by inhibitors of apoptosis (ZVAD), autophagy (chloroquine), or necroptosis (Nec-1) (**Fig. 2h**). LRP8 depletion sensitized cells to hydrogen peroxide (**Fig. 2i**), but not to cell death triggered by the apoptosis-inducer etoposide (**Fig. 2i and Extended Data Fig. 2m**). Consistent with an increased sensitivity to ferroptosis, LRP8KO cells exhibited a significant increase in BODIPY-C11, a reporter of lipid peroxidation, following RSL3 treatment (**Fig. 2j,k**).

**Fig. 3:**
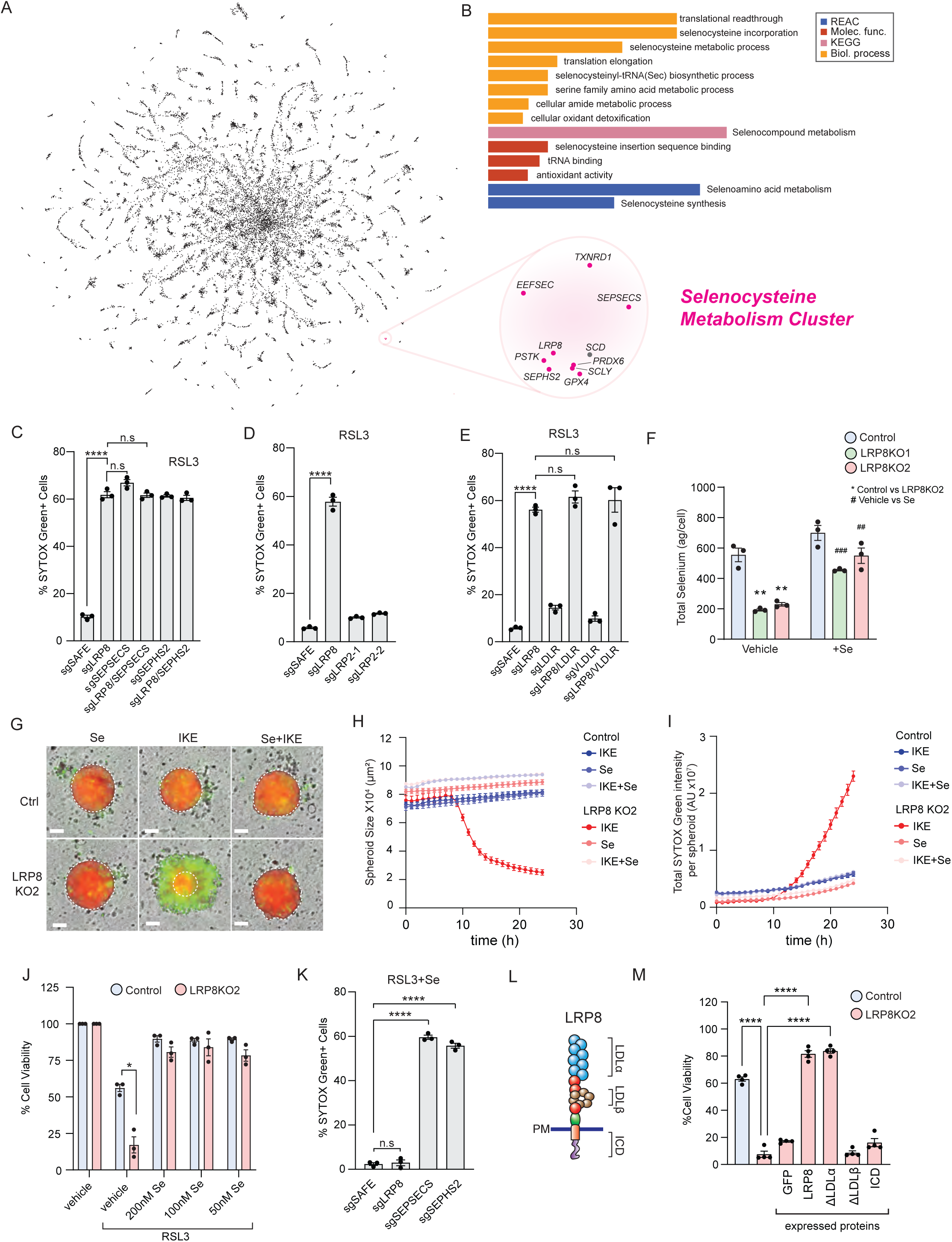
LRP8 regulation of selenium levels impacts ferroptosis resistance. **a**, Overview of genome-wide gene coessentiality, highlighting LRP8 in a selenocysteine metabolism cluster. **b**, Gene ontology (GO) analysis of the genes in the selenocysteine metabolism cluster. **c-e**, Quantification of the percentage of SG+ cells (dead cells) in control and single or double KO cell lines targeting the indicated genes related to selenocysteine metabolism (**c**), lipoprotein receptors (**d**), or LRP2 (**e**). **f**, ICP-MS measurement of the selenium levels in control and LRP8KO cells alone or in the presence of 200 nM Se for 24 hr. **g**, Live-cell imaging of control and LRP8KO spheroids cultured with IKE or Se. Live cells stably express mCherry and dead cells are labeled with SG. **h,i**, Quantification of the size of each spheroid (**h**) or the total intensity of green signal (**i**) over 24 hr. **j**, Cell viability of control and LRP8KO cells upon treatment with 100 nM RSL3 combined with vehicle control or different [Se] for 24 hr. **k**, Quantification of SG+ cells in the indicated KO cells treated with 100 nM RSL3 for 24 hr. **l**, Schematic illustration of LRP8 domains. **m**, Cell viability of control cells or LRP8KO stably expressing the indicated LRP8 mutants in the presence of 100 nM RSL3 for 24 hr. Data represent mean ± S.E.M. of three biological replicates, except for **m**, where n=4. Data were compared using an unpaired t-test (**c-f**) and one-way ANOVA (**m**). In **h** and **i**, at least 8 independent spheroids were collected from three biological repeats.

Ferroptosis can be suppressed by high cell densities and intercellular interactions^16, 17^. Thus, it is crucial to examine whether the effects in 2-dimensional (2-D) cell cultures translate into 3-dimensional (3-D) tumor models, which recapitulate important aspects of *in vivo* tumor biology such as cell-cell interactions, cell interactions with the extracellular matrix, organization of cell layers, cell proliferation, and cell death (e.g. ferroptosis)^18, 19^. The loss of LRP8 had no effect on spheroid growth but dramatically sensitized spheroids to ferroptosis triggered by IKE relative to control cells (**Fig. 2l-n**). The IKE-treated LRP8KO spheroids shrank over time and were surrounded by dead cells labeled by SYTOX Green (**Fig. 2l-n**). The addition of Fer1 blocked cell death and spheroid shrinkage (**Fig. 2l-n**). Together, these findings demonstrate that LRP8 plays a broad role in suppressing ferroptosis in both 2-D and 3-D models of cancer cell growth and survival.

### LRP8 regulation of cellular selenium uptake influences ferroptosis sensitivity

LRP8 has multiple functions that are mediated by its interactions with different ligands, including roles as an ApoE lipoprotein receptor^20^, as a reelin receptor in the central nervous system^21–23^, and as a receptor for the binding and endocytic uptake of the SEC-rich protein selenoprotein P (SEPP1)^7, 24, 25^. To determine how LRP8 might be connected with cell death in cancer cells, we exploited coessentiality tools that take advantage of data from hundreds of CRISPR viability screens to cluster genes with similar profiles of essentiality, enabling functional prediction^26, 27^. These unbiased bioinformatic analyses indicate a strong connection between LRP8 with glutathione metabolism and selenium-related factors (**Fig. 3a,b and Extended Data Fig. 4a**). Indeed, LRP8 exhibited associations with SEC synthesis and degradation enzymes, such as SEPSECS, PSTK, EEFSEC, SEPHS2, and SCLY, and with selenoproteins, such as GPX4 and TXNRD1. Consistent with a model in which LRP8 impacts ferroptosis in cancer cells through regulation of selenium metabolism, CRISPR-mediated depletion of key SEC translation factors (e.g. SEPSECS, SEPHS2, and PSTK) (**Extended Data Fig. 4b**) phenocopied the LRP8KO ferroptosis sensitivity (**Fig. 3c and Extended Data Fig. 4c**). Importantly, double KO of SEPSECS or SEPHS2 with LRP8 had no additive effect on the sensitivity to RSL3 (**Fig. 3c**) or Erastin2 (**Extended Data Fig. 4d**), suggesting that these genes act within the same pathway. While LRP2 has also been implicated in SEPP1 binding and selenium uptake in certain tissues such as the kidney^28^, it was not significantly enriched or disenriched in our screen (**Fig. 1e,f and Supplementary Table S2**) and loss of LRP2 had no effect on ferroptosis sensitivity (**Fig. 3d and Extended Data Fig. 4e**), suggesting that ferroptosis resistance is a specific role for LRP8 in these cells. Depletion of other members of the lipoprotein receptor superfamily, such as the well-characterized lipoprotein receptors VLDR and LDLR, had no impact on ferroptosis resistance (**Fig. 3e and Extended Data Fig. 4e**) and no additional effect was observed when VLDR or LDLR were knocked out together with LRP8 (**Fig. 3e and Extended Data Fig. 4f**).

**Fig. 4:**
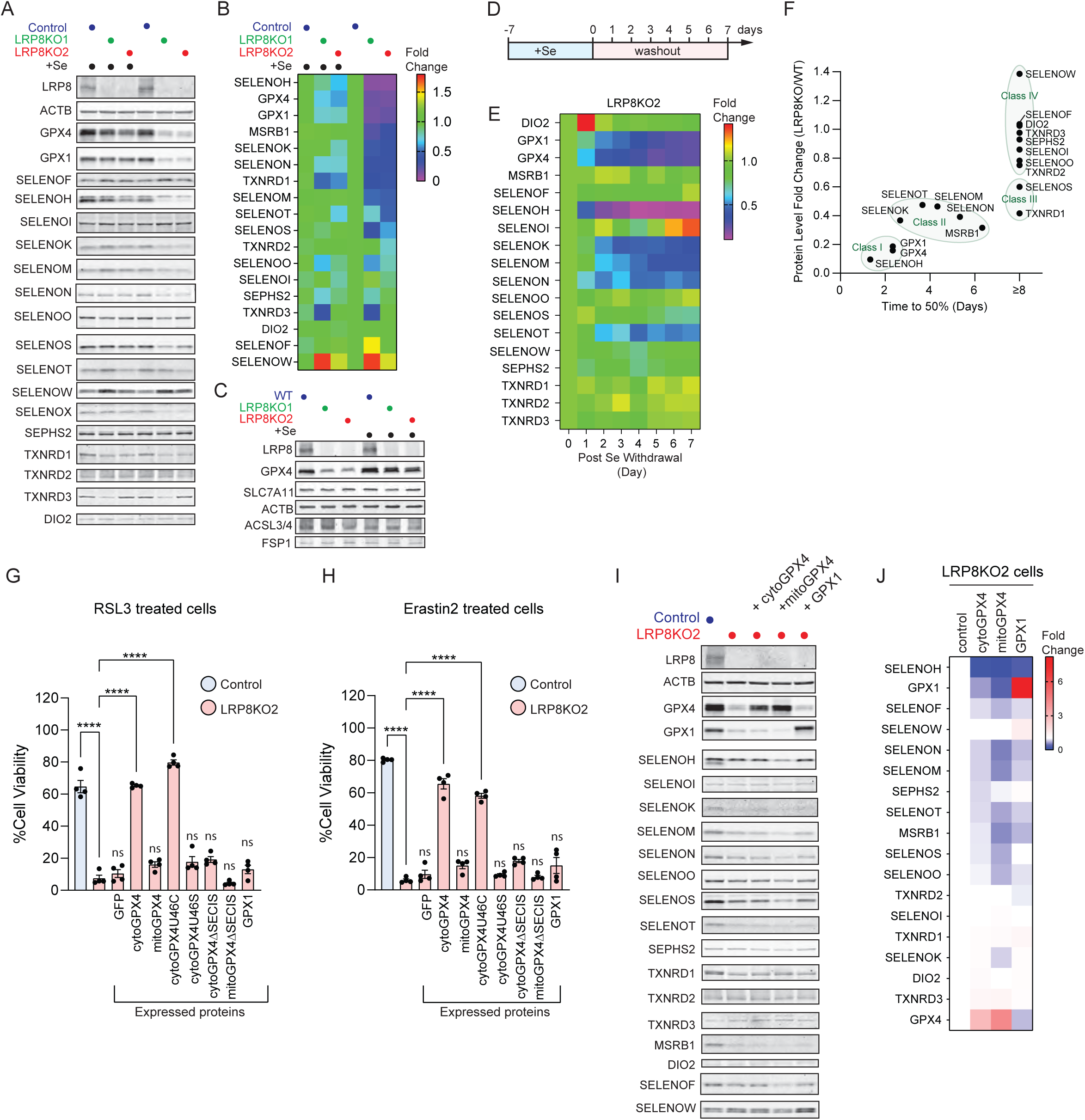
Altered selenocysteine protein expression in LRP8KO cells confers a ferroptosis vulnerability. **a**, Representative immunoblotting of 18 selenocysteine proteins in control and LRP8KO cells. Cell lysates were collected under basal conditions or pre-treated with 200 nM Se for 48 hr. **b**, Quantification of protein levels from panel **a** (*n* = 3 independent experiments). The level of each protein was normalized to the levels of ACTB from the same lysate sample and subsequently normalized to that of control under basal conditions. The fold changes are represented as a heat map. **c**, Western blot analysis of cell lysates from control and LRP8KO cells comparing the protein levels of SLC7A11, ACSL3, ACSL4, and FSP1. **d**, Timeline of the Se washout assay. **e**, Quantification of selenocysteine protein level upon Se washout (*n* = 3 independent experiments). The level of each protein was first normalized to that of ACTB from the same lysate sample and subsequently normalized to the level in the LRP8KO at Day 0. The fold changes are represented as a heat map. **f**, comparison of selenocysteine protein level between the steady state condition (**a,b**) and time to reach 50% of Day 0 upon Se washout (**e**). Selenoproteins that never reached 50% within 7 days of Se withdrawal are indicated as “≥8 Day”. Groups of selenoproteins that responded similarly are indicated by the circles. **g**,**h,** Cell viability of control and LRP8 cells stably expressing the indicated proteins in the presence of 100 nM RSL3 (**g**) or 3 μM Erastin2 (**h**) for 24 hr. **i**, Representative Western blot showing the changes in selenocysteine protein levels upon the overexpression of GPX4 (long or short isoform) or GPX1. **j**, Quantification of protein levels from **i** (*n* = 3 independent experiments). The level of each protein was first normalized to that of ACTB from the same lysate sample and subsequently normalized to that of LRP8KO. The fold changes were represented as a heat map. Data represent mean ± S.E.M. of three biological replicates.

Measurement of selenium levels using inductively coupled plasma mass spectrometry (ICP-MS) revealed a ∼60% decrease in selenium levels in LRP8KO cells (**Fig. 3f**) , with no change in other elements such as iron, copper, manganese, and zinc (**Extended Data Fig. 5a-d**) nor any change in glutathione levels (**Extended Data Fig. 5e**). The addition of selenite (Se), a form of selenium taken up by cells independent of LRP8, was sufficient to correct selenium levels (**Fig. 3f**) and rescue LRP8KO ferroptosis resistance in 3-D spheroids (**Fig. 3g-i**) and 2-D cultures (**Fig. 3j**). As expected, Se treatment did not impact RSL3- or Erastin2-induced ferroptosis in the cells lacking key enzymes in the synthesis of SEC, SEPSECS or SEPHS2 deficient cells (**Fig. 3k and Extended Data Fig. 5f**). These data demonstrate that the protective activity of Se requires selenoprotein synthesis and is downstream of LRP8. Expression of GFP-tagged full-length LRP8 and LRP8ΔLDLα, which is unable to bind lipoproteins^29^, rescued LRP8KO cells from ferroptosis (**Fig. 3l,m and Extended Data 5g,h**). In contrast, LRP8ΔLDLβ, which is unable to bind SEPP1^30^, was unable to rescue ferroptosis resistance (**Fig. 3l,m and Extended Data 5g,h**). Together, these data indicate that LRP8 promotes ferroptosis resistance by maintaining cellular pools of selenium through SEPP1 binding and uptake.

**Fig. 5:**
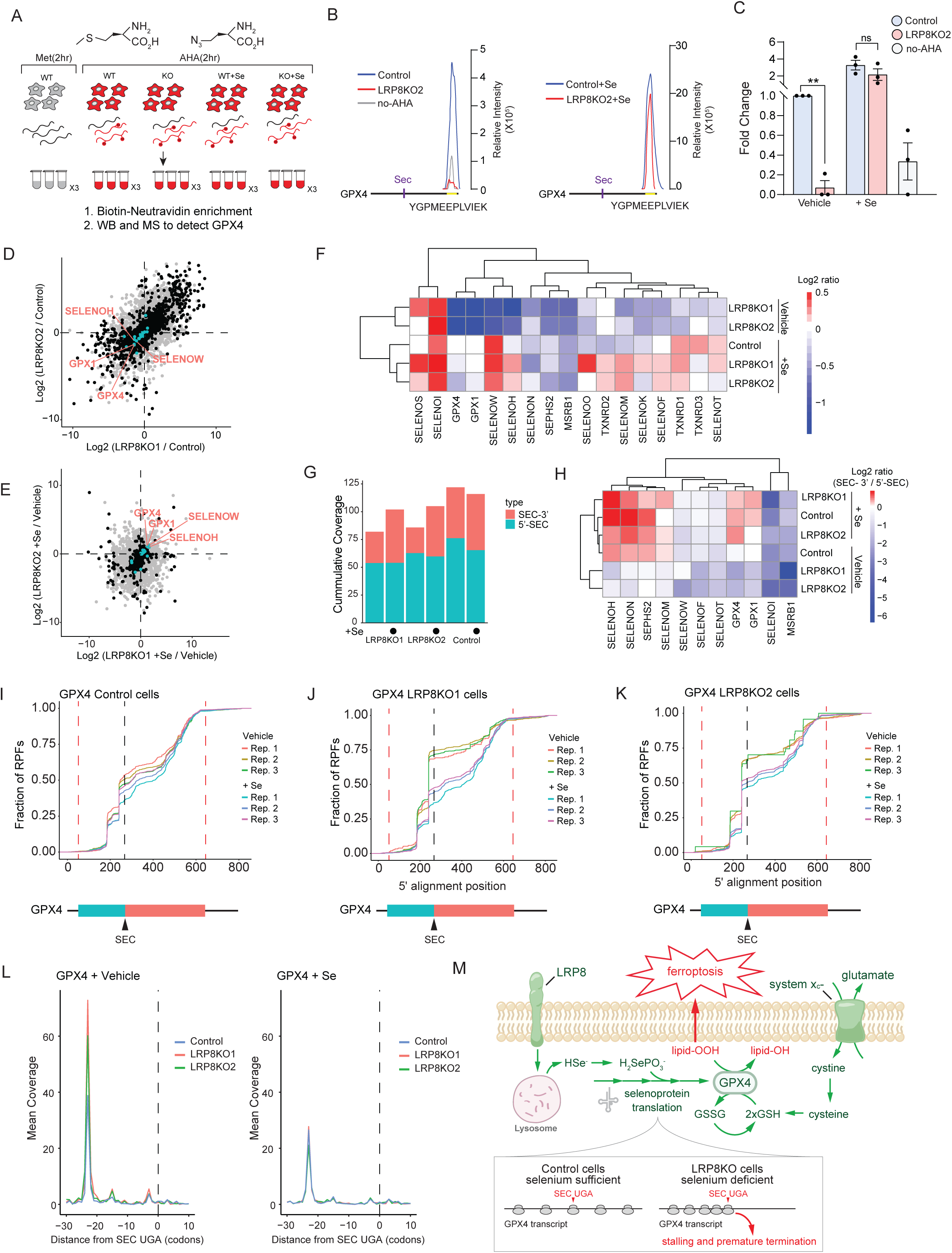
GPX4 is susceptible to ribosome stalling and early translation termination under selenium-limiting conditions. **a**, Experimental workflow for AHA translational analysis of GPX4 in control and LRP8KO cells with or without Se treatment. **b**, Representative LC traces of a GPX4 peptide from AHA labelled control, LRP8KO, and no-AHA cell lysates. **c**, Quantification of the relative abundance of the GPX4 peptide from **b** (*n* = 3 biological replicates). The relative abundance of each sample was first measured by relative intensity and normalized to the level of AHA labeled control from the same replicate. **d**, Scatter plots of fold changes in translation efficiency (TE) between control and two LRP8KOs, respectively. Transcripts with significant TE changes are labeled as black dots. All selenoprotein transcripts are highlighted in cyan. **e**, Fold changes in TE for the two LRP8KO cells in response to Se treatment. **f**, Heatmap summary from panels (**d**) and (**e**) showing the TE fold change for 17 selenoproteins in the indicated cell lines in the presence and absence of Se treatment. The TE from all groups was normalized to the TE from control cells under steady-state (vehicle) and represented as log2 fold changes. **g**, Quantification of the overall density of ribosome-protected fragments (RPFs) from 5’ (Cyan) and 3’ (Red) of the SEC in control and LRP8KOs under steady-state or Se-treated conditions. **h**, Heat map summary of the fold change in the ratio of RPFs 3’ / 5’ of the SEC codon from selected selenoprotein gene transcripts. Those with low counts or a short distance between the SEC and stop codon (<10 nt) were excluded. **i-k**, Empirical Cumulative Distribution Function of GPX4 RPFs in control (**i**), LRP8KO1 (**j**), and LRP8KO2 (**k**) cells incubated with vehicle or Se. The positions of the start and stop codon are indicated by red dash lines, and the location of SEC is marked by a black dash line and black arrowhead. **l**, RPFs accumulated at the 5’ position of the SEC locus on the GPX4 transcript under steady-state (left) or following Se treatment (right). **m**, A model illustrating the mechanism by which LRP8 suppresses ferroptosis. LRP8-mediates the uptake of the SEC-rich protein SEPP1, which is broken down in the lysosome to release selenium. Selenium is employed in the translation of selenoproteins such as GPX4. In the absence of LRP8, selenium becomes limiting for the translation of a subset of selenoproteins, including GPX4. Prolonged ribosome stalling at the GPX4 SEC UGA causes ribosome collisions and early translation termination. The resulting decrease in GPX4 levels sensitizes cells to ferroptosis induction by agents targeting the glutathione-GPX4 pathway.

### LRP8 impacts ferroptosis resistance by promoting GPX4 expression

To test the hypothesis that selenium pools become limiting for the biosynthesis of selenoproteins in the LRP8KO cells, we examined the levels of 18 selenoproteins expressed in HCC1143 TNBC cells. 10 of the 18 selenoproteins were reduced in LRP8KO cells (**Fig. 4a,b and Extended Data Fig. 6**), including the well-known selenium responsive protein GPX1^7, 8^, which is reduced to provide selenium for the biosynthesis of essential selenoproteins. In line with the cellular importance of GPX4, the expression of GPX4 is normally maintained in most tissues during periods of selenium depletion and GPX4 is widely considered to be a constitutively expressed housekeeping protein^1, 7, 8^. Unexpectedly, GPX4 levels were severely reduced in the LRP8KO cells (**Fig. 4a,b and Extended Data Fig. 6**). The levels of GPX4, and other reduced selenoproteins, were rescued by the addition of Se (**Fig. 4a,b and Extended Data Fig. 6**) but not by ferroptosis inhibitors (**Extended Data Fig. 7a**). We observed a similar reduction in GPX4 levels in MDA-MB-453, HCC1937, U87-MG, and SK-MEL-28 LRP8KO cells (**Extended Data Fig. 7b-f**). Importantly, the loss of GPX4 was reversed by re-expressing full-length LRP8 and LRP8ΔLDLα, but not LRP8ΔLDLβ or LRP8ICD (LRP8 intracellular domain) which are unable to bind SEPP1 (**Extended Data Fig. 7g,h**). LRP8KO did not affect the levels of other known ferroptosis regulators that do not contain SEC, such as ACSL3, ACSL4, FSP1, and SLC7A11 (**Fig. 4c**).

To further assess the dynamic changes in selenoprotein levels under selenium limiting conditions, we pre-treated LRP8KO cells with Se for seven days to restore the selenoprotein levels and then quantified selenoprotein levels following Se removal (**Fig 4d,e and Extended Data Fig. 8a,b**). Analysis of selenoprotein levels under steady-state conditions and their rate of change following Se washout revealed that the 18 selenoproteins could be generally grouped into distinct classes (**Fig. 4f**). Class I selenoproteins, which included GPX4, GPX1, and SELENOH, exhibited the largest reduction in levels under basal conditions and were rapidly lost following Se washout, decreasing to 50% within ∼1.3, 2.3, and 2.3 days of selenium withdrawal, respectively (**Fig. 4f**). Contrary to the current dogma of the selenoprotein hierarchy^7, 8^, our data reveal that GPX4 is one of the first proteins to be reduced following selenium depletion in cancer cells.

To address whether the decrease in GPX4 is responsible for the ferroptosis sensitivity in LRP8KO cells, we measured the effect of overexpressing GPX4. Expression of the cytosolic short isoform of GPX4 (cytoGPX4) (**Extended Data Fig. 9a,b**), but not the mitochondrial long isoform of GPX4 (mitoGPX4) (**Extended Data Fig. 9a,b**), rescued ferroptosis resistance in LRP8KO cells (**Fig. 4g,h**). In addition, expression of a mutant cytoGPX4 in which the SEC was replaced by cysteine (cytoGPX4U46C), but not a mutant in which the SEC was replaced by serine (cytoGPX4U46S), also restored ferroptosis resistance of LRP8KO cells in the presence of RSL3 or Erastin2 (**Fig. 4g,h and Extended Data Fig. 9a**). These data are consistent with previous findings that cytoGPX4U46C retains partial activity^5^. As expected, GPX4 constructs lacking the selenocysteine insertion sequence (SECIS) (mitoGPX4ΔSECIS and cytoGPX4ΔSECIS) were not expressed and failed to rescue ferroptosis resistance (**Fig. 4g,h and Extended Data Fig. 9a**). In contrast to GPX4, overexpression of GPX1 in LRP8KO cells had no effect on RSL3- or Erastin2-induced ferroptosis (**Fig. 4g,h**). The overexpression of cytoGPX4, mitoGPX4, and GPX1 resulted in small additional decrease in the selenium- responsive selenoproteins, but had no effect on the levels of the well maintained Class IV selenoproteins (**Fig. 4i,j and Extended Data Fig. 10**). These data are consistent with a limiting pool of available selenium that influences selenoprotein levels in the LRP8KO cells. Together, these findings demonstrate that LRP8 promotes ferroptosis by supporting the selenium-dependent expression of GPX4. In addition, in contrast to many tissues where GPX4 is maintained^7, 8, 31^, GPX4 levels decrease when selenium is limiting in cancer cells, indicating a cancer-specific ferroptosis vulnerability.

### Loss of selenium disrupts GPX4 translation

Selenoproteins are subject to complex regulation, including hierarchical transcriptional and posttranscriptional mechanisms that adapt selenoprotein expression to fluctuations in selenium levels^7^. Given the critical function of GPX4 and its generally accepted position at the top of the selenoprotein hierarchy^7, 8^, we were surprised by the reduction in GPX4 levels in the selenium- deficient LRP8KO cells and thus sought to understand the underlying mechanism of GPX4 regulation. Both RNAseq and qRT-PCR data indicated that GPX4 transcript levels were unaltered in LRP8KO cell lines and were unaffected by Se treatment (**Extended Data Fig. 11a and Supplementary Table 3**). We also examined whether GPX4 is targeted for degradation, but inhibiting the proteasome with MG132 and lysosome with chloroquine had no effect on GPX4 levels (**Extended Data Fig. 11b**).

To investigate if the translation of GPX4 is disturbed in LRP8KO cells, we employed BioOrthogonal Non-Canonical Amino acid Tagging (BONCAT) together with proteomics and western blotting. Cells were metabolically labeled with a methionine analog L-azidohomoalanine (AHA) to tag newly synthesized proteins (**Fig. 5a**). Newly synthesized, AHA-labeled proteins were conjugated to a biotin moiety using click chemistry and then affinity purified for analysis. By mass-spectrometry, we detected two GPX4 peptides. The major peptide (YGPMEEPLVIEK) resided at the end of the protein (153-164) and was captured in almost all samples. Using this peptide to represent the level of newly synthesized GPX4, we observed a substantially larger peak for control cells relative to LRP8KO cells (**Fig. 5b,c**). Se treatment drastically diminished the difference in GPX4 levels between control and LRP8KO cells (**Fig. 5b,c)**. Similar results were obtained by western blotting (**Extended Data Fig. 11c,d**), indicating that the limiting selenium in LRP8KO cells attenuates translation of full length GPX4.

### Ribosome stalling and early translation termination for GPX4

To systematically examine protein translation in LRP8KO cells, we employed ribosome profiling, an approach that uses deep sequencing of ribosome-protected fragments (RPFs)^32, 33^. Normalization of RPFs to mRNA abundance data from RNA-seq yields a measure of translation efficiency (TE). These analyses revealed a decrease in the TE for several of the selenoproteins in both LRP8KO cell lines, including GPX4, GPX1, SELENOH, and SELENOW (**Fig. 5d-f, Extended Data Fig. 12, and Supplementary Table 4**). The pattern of RPFs across the selenoprotein transcripts demonstrated an altered distribution in the LRP8KO cells, with a general decrease in RPFs downstream, 3’ of the SEC UGA relative to the upstream, 5’ portion of the reading frame (**Fig. 5g,h**). The reduction in downstream RPFs was particularly strong for a subset of selenoproteins, including GPX4 (**Fig. 5g,h**). Notably, the decrease in GPX4 TE (**Fig. 5d-f**) and downstream RPFs were both reversed by the addition of Se (**Fig. 5h**).

The density of ribosome footprints positioned at a specific codon indicates the average ribosome dwell time and thus reflects relative ribosome translation speed across that codon^32, 33^. Several selenoprotein transcripts exhibited an accumulation of ribosomes (i.e. a ribosome pause or stall) just prior to the SEC UGA (**Extended Data Fig. 13**). GPX4 showed evidence of stalled ribosomes in control cells (**Fig. 5i**), consistent with slow decoding of the SEC UGA even under basal conditions. However, ribosomes staling on GPX4 was severely exacerbated in the LRP8KO cells, with ∼50% of the RPFs positioned just prior to the SEC UGA (**Fig. 5j,k**). The increase in stalled ribosomes in the LRP8KOs was largely rescued by Se addition (**Fig. 5j,k**). Together with the decrease in RPFs 3’ of the SEC UGA (**Fig.5g,h**), these data suggest that the reduced GPX4 TE is due to ribosome stalling at the SEC UGA and subsequent ribosome dissociation (i.e. early termination).

Stalled translation can lead to collisions between ribosomes and accumulation of tightly apposed poly-ribosome clusters^34, 35^. While some selenoprotein transcripts, such as SELENOH, showed footprints from single ribosome (i.e. monosomes) stalled while decoding the SEC UGA (**Extended Data Fig. 14a**). However, for other selenoprotein transcripts, the ribosomes were stalled further upstream of the SEC UGA (**Extended Data Fig. 14**). These upstream stalls were exacerbated in Se-deficient LRP8KO cells, suggesting that they result from slow SEC decoding that cause collisions with trailing ribosomes.The upstream position of the stalled ribosome and the distance from the SEC UGA is consistent with stacked ribosomes, with two collided ribosomes (i.e. a disome) for GPX1, SELENOT, and SELENOM (**Extended Data Fig. 14b-d)** and three collided ribosomes (i.e. a trisome) evident for GPX4 (**Fig. 5l**). The trisome peak upstream of the GPX4 SEC UGA was highly pronounced in the LRP8KO cells (**Fig. 5l**) and substantially reduced by Se treatment (**Fig. 5l**). Together, these data implicate ribosome collisions and early translation termination due to ribosome stalling as the cause of the low GPX4 levels in the selenium-deficient LRP8KO cells.

In this study, we identify LRP8 as a ferroptosis resistance factor that promotes GPX4 translation through its role in regulating cellular selenium levels (**Fig. 5m**). LRP8 is a promising candidate therapeutic target that is upregulated in cancer, positively correlated with ferroptosis resistance across hundreds of cancer cell lines, and has a limited tissue expression profile, being most highly expressed in the brain and testes^20^. Consistent with its potential as a therapeutic target, our data demonstrate that loss of LRP8 is synthetic lethal with ferroptosis inducers in 2-D and 3- D cancer cell models, and previous findings indicate that LRP8KO reduces tumor growth in xenografts^36, 37^. Our results support a model in which the translation of full-length GPX4 is reduced in LRP8KO cells due to limiting selenium (**Fig. 5m**). Remarkably, the decrease in GPX4 translation is not part of the known active hierarchical regulation of selenoproteins^7^, but is instead due to the propensity of ribosomes to stall at the GPX4 SEC UGA. The prolonged ribosome stalling in the selenium deficient LRP8KO cells imparts a susceptibily for catastrophic ribosome collisions and early translation termination (**Fig. 5m**). Ribosome collisions are known to trigger a ribosome quality control pathway that mediates ribosome dissociation and proteasomal clearance of ribosome-associated protein fragments^38, 39^. While the high amount of GPX4 translation is likely to be a protective cancer adaptation that promotes resistance to oxidative lipid damage, our data demonstrate that it becomes a liability during periods of low selenium. Together, our findings reveal ribosome stalling, ribosome collisions, and early GPX4 translation termination via disruptions in selenium metabolism as targetable vulnerabilities that sensitize cancer cells to ferroptosis.

## ACKNOWLEDGEMENTS

This research was supported by grants from the National Institutes of Health (R01GM112948 to J.A.O., T32GM007232 to L.F., F31DK121477 to M.A.R., 1R01GM122923 to S.J.D.), American Cancer Society (Research Scholar Award RSG-19-192-01 to J.A.O.), and Melanoma Research Alliance (Award 620458 to J.A.O.). J.A.O. is a Chan Zuckerberg Biohub investigator and Miller Institute Professor. We thank Drs. Mike Lange and Kirill Bersuker for critical reading of the manuscript. ICP-MS measurements were performed in the OHSU Elemental Analysis Core with partial support from NIH (S10RR025512). We thank Dr. Steve Eyles (UMass Amherst, RRID: SCR_019063) for assistance with high-resolution MS acquired on an Orbitrap Fusion mass spectrometer funded by NIH grant 1S10OD010645-01A1.

## AUTHOR CONTRIBUTIONS

Z.L. and J.A.O. conceived of the project and designed the experiments. J.A.O. and Z.I. wrote the majority of the manuscript. All authors read, edited, and contributed to the manuscript. Z.L. performed the majority of the experiments. Z.L. and L.F. prepared samples for ribosome profiling, and L.F. and N.T.I. analyzed the sequencing data. Z.L. and K.K.D. performed and analyzed protein translation using AHA-labeling and proteomics. Z.L. performed and analyzed the CRISPR screen with assistance from M.A.R. and M.C.B. S.J.D. and L.M. performed the glutathione measurements.

## AUTHOR INFORMATION

Correspondence and requests for materials should be addressed to J.A.O.

## COMPETING INTERESTS

S.J.D. is a member of the scientific advisory board for Ferro Therapeutics.

## METHODS

### Cell Lines and culture conditions

U-2 OS Tet-On cells were purchased from Clontech (#630919). MDA-MB-453 cells were purchased from the American Type Culture Collection (ATCC, # HTB-131). U-87 MG, A-375, SK-MEL-28, HCC1143, and HCC1937 were all obtained from UC Berkeley Cell Culture Facility. U-2 OS, MDA-MB-453, U-87 MG, A-375, SK-MEL-28, HEK293T were maintained in DMEM containing 4.5 g/l glucose and L-glutamine (Corning). HCC1143 and HCC1937 were cultured in RPMI 1640 with L-glutamine (Gibco). All media were supplemented with 10% fetal bovine serum (FBS, Gemini Bio Products), and all cell lines were grown at 37 °C with 5% CO2. All cell lines were tested for mycoplasma.

### Generation of CRISPR-Cas9 genome-edited cell lines

For the CRISPR–Cas9 synthetic lethal screen, U-2 OS Tet-On and MDA-MB-453 lines stably expressing Cas9 were generated by infection with lentiCas9-Blast, a gift from F. Zhang (Addgene plasmid no. 52962) and cells were selected in medium containing 10μg/ml blasticidin (Gibco, # A1113903). Active Cas9 expression was validated by flow cytometry analysis following infection with a self-cutting mCherry plasmid (a gift from M. Bassik, Stanford University), which expresses mCherry and a sgRNA targeting the mCherry gene.

LRP8KO lines in MDA-MB-453, HCC1143, HCC1937, U-87 MG, A-375, SK-MEL-28 were generated by infection with lentiCas9-Blast and pMCB320 plasmid (gift from M. Bassik, Stanford University) encoding LRP8-targeting sgRNAs. Cells were selected in media containing 10 μg/ml blasticidin and 1 μg/ml puromycin. Single clones of LRP8KO from MDA-MB-453, HCC1143, and HCC1937 were isolated using cloning rings or serial dilution.

### Plasmids

Two CRISPR single guide RNA (sgRNA) sequences targeting LRP8 and one targeting SEPHS2 were selected based on the effective score from the whole human CRISPR knockout library used for genetic screening. The oligonucleotide sequences preceding the protospacer motif were: sgLRP8-1, 5’GCCGGCCAAGGATTGCGAAA3’; sgLRP8-2, 5’ GCTTAGACCACAGCGACG3’; sgSEHPS2, 5’ GGTGTTAACCAAACCGTT3’. One other sgRNA sequence targeting LRP8 and sgRNAs targeting SEPSECS, LDLR, VLDLR and LRP2 were picked from pre-designed sequences by Integrated DNA Technologies (IDT): sgLRP8-3,5’ GCCACTGCATCCACGAACGG3’; sgSEPSECS,5’ GCTCGCATGAGCACCTCATA’; sgLDLR,5’ GACGAGTTTCGCTGCCACGA3’; sgVLDLR,5’ GTCAGGTCGGCAAGTTCGAG3’; sgLRP2-1,5’ CATACGGCGTCTCATGGCAC 3’; sgLRP2-2,5’ AGTGAACTGGTAACCACCGC3’; All guide sequences were inserted into pMCB320 between BstX1 and BlpI.

Wildtype full-length LRP8 was cloned from a MDA-MB-453 complementary DNA (cDNA) library and inserted into pLenti-CMV-GFP-Hygro (Addgene #17446) between XbaI and BamHI. The PAM sequence for LRP8 guide 3 was then mutated without changing the encoded amino acids. LRP8ΔLDLα(Δ46-334), LRP8ΔLDLβ(Δ462-681), LRP8ICD(Δ2-847) were sub-cloned from LRP8-GFP containing the PAM mutation.

Full-length cytoGPX4 and mitoGPX4 containing the SECIS element were synthesized using a gBlock, a synthesized double-stranded DNA fragment from IDT and inserted into pENTR1A-GFP-N2 (Addgene # 19364) between EcoRI and NotI, and GFP was removed. Both were subsequently cloned into pLenti-CMV-Hygro-DEST (Addgene # 17454) by gateway cloning. To make cytoGPX4ΔSECIS-GFP and mitoGPX4ΔSECIS-GFP, cytoGPX4 and mitoGPX4 were cloned from a MDA-MB-453 complementary DNA (cDNA) library and inserted into pENTR1A-GFP-N2 between EcoRI and KpnI, and then cloned into pLenti-CMV-Hygro-DEST. cytoGPX4U46C-GFP and cytoGPX4U46S-GFP were generated by site-directed mutagenesis from cytoGPX4ΔSECIS-GFP. GPX1 containing the SECIS element was synthesized as a gBlock from IDT, inserted into pENTR1A-GFP-N2 between the KpnI and NotI sites, and gateway cloned into pLenti-CMV-Hygro-DEST.

### Chemicals and reagents

1S,3R-RSL3 (referred to as RSL3) (#19288), Ferrostatin-1 (#17729), idebenone (#15475), Deferoxamine (DFO, #14595), Erastin2 (#27087), ML162 (#20455), ZVAD(OMe)-FMK (#27421), Necrostatin-1 (#11658), IKE (Imidazole Ketone Erastin, #27088), Tocopherol (#25985) were all purchased from Cayman Chemical. Etoposide (#E1383), chloroquine (#C6628), and sodium selenite (#S5261), were all purchased from Sigma-Aldrich. Blasticidin (A1113903), Puromycin (A1113803), SYTOX Green Dead Cell Stain (#34860), and MitoTracker Deep Red FM (M22426) were all purchased from Thermo Fisher Scientific.

### Cell viability and death analysis

For cell death analysis using the Essen IncuCyte Live-Cell imaging system (Sartorius), cells were plated at a density of 5,000 cells per well in black 96-well plates (Corning, #3904) the day before imaging. The next day, the media were replaced with fresh media containing 30 nM SYTOX Green Dead Cell Stain (Invitrogen, #S34860) and compounds at different doses. The plates were transferred to an IncuCyte Zoom or S3 Live-Cell imaging system enclosed in an incubator set to 37 °C and 5% CO2. Three images per well were captured in the green and red channels every 1 or 2 hr over a 24 hr period, and the ratio of SYTOX Green-positive objects (dead cells) to mCherry positive objects (live cells) was quantified using the Sartorius image analysis software. For each treatment condition, the SYTOX-to-total (SYTOX+mCherry)-object ratio was plotted against the 24 h imaging interval, the AUC (Area under the curve) was calculated, and the average AUC was plotted as a function of drug concentration (for example, RSL3) using Prism (GraphPad).

For cell viability analysis, the CellTiter-Glo assay was performed according to the manufacturer’s instructions. Briefly, 5,000 cells were seeded in each well of a white 96-well plate (Corning, #3917) the day before compound treatments. The next day, media was replaced with 100 μL fresh medium containing compounds at a different dose. Plates were incubated at 37 °C for 24 hr and then at room temperature for 30 min. 100 μL CellTiter-Glo 2.0 reagent (Promega, #G9242) was mixed with the media. The plates were incubated at room temperature for 10 mins, and the luminescent signal was recorded with a SpectraMax i3 (Molecular Devices).

### Western blotting

Cells were washed twice with phosphate-buffered saline (PBS), lysed in RIPA buffer supplemented with 1x Pierce EDTA-free Protease Inhibitor Cocktail (Thermo Scientific #A32963), and sonicated for 15 sec. The cell lysate was then centrifuged for 10 min at 15,000*g* to remove any cell debris. Protein concentrations were determined using the bicinchoninic acid (BCA) protein assay (Thermo Fisher Scientific), and equal amounts of protein by weight were combined with Laemmli buffer, boiled for 5 min at 95 °C, separated on 4–20% polyacrylamide gradient gels (Bio-Rad Laboratories), and transferred onto nitrocellulose membranes (Bio-Rad Laboratories). Membranes were blocked in PBS-containing 0.1% Tween 20 (PBST) containing 5% (w/v) dried milk for 60 min. Membranes were incubated overnight at 4 °C in PBST containing 3% bovine serum albumin (BSA) (Sigma Aldrich) and primary antibodies. After 3 washes with PBST, membranes were incubated at room temperature for 60 min in 3% BSA in PBST containing fluorescent secondary antibodies, followed by 3 more washes with PBST. Immunoblots were then imaged on an LI-COR imager (LI-COR Biosciences).

The following blotting reagents and antibodies were used: GPX4 (Abcam, #ab125066), GPX4 (Invitrogen, #LF-MA0085), GPX1 (Abcam, #ab22604), SEPHS2 (Sigma-Aldrich, #SAB2700271), SELENOH (Sigma-Aldrich, #HPA048362), SELENOF (Sigma-Aldrich, #HPA054937), SELENOS (Sigma-Aldrich, #HPA010025), SELENOT (Sigma-Aldrich, #HPA039780), SELENOK (Sigma-Aldrich, #HPA008196), MSRB1 (Proteintech, #15333-1-AP), SELENOI (Novus Biologicals, # H00085465-A01), SELENOO (Abcam, # ab172957), SELENOM (Santa Cruz, #sc-514952), SELENON (Santa Cruz, #sc-365824), SELENOW (Rockland,# 600-401-A29S), TXNRD1 (Santa Cruz, #sc-28321), TXNRD2 (Proteintech, # 16360-1-AP), TXNRD3 (Proteintech, #19517-1-AP), DIO2 (Proteintech, #26513-1-AP), LRP8 (Abcam, #ab108208), ACTB (Santa Cruz, #sc-47778), α-Tubulin (Cell Signaling, #2144), FSP1 (Santa Cruz, #sc-377120), SLC7A11 (Cell signaling, #12691), ACSL3/ACSL4 (Sigma-Aldrich, #SAB2701949), GFP (Santa Cruz, #sc-9996) IRDye800 conjugated secondary (LI-COR Biosciences, #926-32211) and anti-mouse Alexa Fluor 680 conjugated secondary (Invitrogen, #A21058).

### Fluorescence microscopy

Cells were washed 3x with Dulbecco’s phosphate-buffered saline (DPBS), fixed for 20 min in DPBS containing 4% (w/v) paraformaldehyde, and washed 3x again with DPBS. Cells were permeabilized for 2 min with DPBS containing 0.2% Triton X-100, washed 3x with DPBS, and incubated in blocking solution (1% BSA, 0.3M glycine in DPBS) for 30 min. Cells were incubated with anti-GPX4 antibody in blocking solution (1:100 dilution) overnight at 4 °C, washed 3x, and incubated for 1 h at room temperature in blocking solution containing anti-rabbit secondary antibody conjugated to Alexa Fluor 488 (1:250 dilution) (Thermo Fisher Scientific). After an additional 3x washes, coverslips were mounted on glass slides using Permount mounting medium (Fisher Chemical). Images were acquired using an epifluorescence microscope (Axio Observer 7, Carl Zeiss) fitted with a 63x plan-Apochromat oil objective [Numerical Aperture (NA),1.4]. Digital images were captured with cameras (Axiocam 712 mono, Zeiss) using Zeiss Zen 3.2 software.

### CRISPR–Cas9 synthetic lethal screen

The CRISPR–Cas9 screen was performed as previously described^9^. The genome-wide human CRISPR sgRNA knockout library contains 9 sublibraries, comprising in total 225,171 elements—including 212,821 sgRNAs targeting 20,549 genes (about 10 sgRNAs per gene) and 12,350 negative-control sgRNAs. To generate lentiviral particles, each sublibrary was co-transfected with third-generation lentiviral packaging plasmids (pVSVG, pRSV and pMDL) into HEK293T cells. Media containing lentivirus was collected 48 and 72 h after transfection, combined, filtered, and then used to infect approximately 2.0 × 10^8^ U-2 OS Tet-On cells or MDA-MB-453 cells stably expressing Cas9. After 72 hr of growth, infected cells were selected in media containing 1 μg/ml puromycin until over 90% cells were mCherry-positive. Cells were then re-seeded in 500-cm^2^ plates (about 8 × 10^6^ cells per plate) and recovered in media lacking puromycin for 24 h. For the screen, a total of about 2.5 ×10^8^ cells (∼1,000-fold library coverage) were treated with either DMSO or 0.5 μM RSL3 for 10 days. Cells were then trypsinized, collected by centrifugation at 1,000*g*, washed twice with PBS, and pellets were frozen at -80 °C. Genomic DNA was extracted using the QIAamp DNA Blood Maxi Kit (QIAGEN) according to the manufacturer’s instructions. sgRNA sequence libraries were prepared from genomic DNA by two rounds of PCR using the Herculase II Fusion DNA Polymerase (Agilent). sgRNA sequences were amplified by the primers oMCB1562 and oMCB1563 and then indexed using the Illumina TruSeq LT adaptor sequences (AD002, AD004, AD006, AD009, AD0012, AD0014, AD0016, AD0019, AD0022 for DMSO and AD003, AD005, AD007, AD010, AD0013, AD0015, AD0018, AD0021, AD0025 for RSL3), for downstream deep sequencing analysis. PCR products were separated on a 2% tris-borate-EDTA (TBE)-agarose gel, purified using the QIAquick Gel Extraction Kit (Qiagen) and assessed for quality using a Fragment Analyzer (Agilent). PCR amplicons from each sample were pooled in a 1:1 ratio (DMSO: RSL3) based on their concentrations as determined by Qubit Fluorometric Quantification. Then each sample-pair was pooled together in a ratio based on the number of elements in each sublibrary. sgRNA sequences were analyzed by deep sequencing using the primer oMCB1672 on an Illumina NextSeq instrument at the Oklahoma Medical Research Foundation. Sequence reads were aligned to the sgRNA reference library using Bowtie software. For each gene, a gene effect and score (likely maximum effect size and score), and *p*-value were calculated using the casTLE statistical framework as previously described^40, 41^.

### Competitive growth assay

5 x 10^4^ WT (no mCherry tag) and sgLRP8 (with mCherry tag) cells were mixed together and seeded in wells of a 6-well plate. 48 hr after the seeding, cells were treated with various compounds and cultured at 37 °C for another 48 hr. Live cells were then collected and analyzed by flow cytometry on a BD LSRFortessa to measure the ratio of mCherry negative to mCherry positive cells. Data were analyzed using FlowJo.

### BODIPY 581/591 C11 analysis

Cells seeded in a 6-well plate were treated with 200nM RSL3 for 5 hr and washed once with DPBS containing calcium and magnesium. Cells then were incubated in DPBS containing 5μM BODIPY 581/591 C11 (Invitrogen, #D3861) at 37 °C for 10 min and washed 3x with DPBS without calcium or magnesium. Cells were detached from the plate with trypsin, and green fluorescence was analyzed by flow cytometry (>10000 cells) on a BD LSRFortessa. Data were analyzed using FlowJo.

### Selenium measurement by ICP-MS

Inductively coupled plasma mass spectrometry (ICP-MS) measurements were performed at the OHSU Elemental Analysis Core. Cells were harvested directly into 15 ml metal-free centrifuge tubes (VWR, #89049-170). An aliquot was removed for cell counting, and then media was removed (as much as possible). For digestion, 100 µl of concentrated HNO_3_ (trace metal grade, Fisher # 7697-37-2) was added to each cell pellet (containing approximately 10^6^ cells), the tubes were loosely capped, and the samples were heated for 1-2 hrs at 90°C on a heating block. 1500 - 2000 µl of 1% HNO_3_ was added to each sample after digestion. The following digestion and background controls were prepared (each in triplicate): (1) NIST bovine liver standard (SRM 1577c): ∼10 mg (weighed on a precision balance (+/- 0.1 mg)) was added to 200 µl of 50% HNO_3_ (prepared from trace metal grade HNO_3_) and heated for 2 hrs at 90°C in a loosely capped 15 ml centrifuge tube (metal free, VWR, #490001), (2) Solution standard: 2 µl of calibration standard #2 (“CEM2”, LGC Standards #VHG-SM70B-100) was added to 200 µl of 50% HNO_3_ and heated for 2 hrs at 90°C in a loosely capped 15ml centrifuge tube (VWR, #490001), (3) Blank: 200 µl of 50% HNO_3_ (prepared from trace metal grade HNO_3_) was added to an empty tube and heated for 2 hr at 90°C in a loosely capped 15ml centrifuge tube (VWR, #490001).

ICP-MS analysis was performed using an Agilent 7700x equipped with an ASX 500 autosampler. The system was operated at a radio frequency power of 1550 W, an argon plasma gas flow rate of 15 L/min, and an Ar carrier gas flow rate of 0.9 L/min. Elements were measured in either kinetic energy discrimination (KED) mode using He gas (4.3 ml/min) (Mn, Fe, Cu, Zn) or in mass-on-mass (hydrogen) mode using H_2_ gas (3.4 ml/min) (Se). Data were quantified using weighed, serial dilutions of a multi-element standard (“CEM 2”, (VHG-SM70B-100) Mn, Fe, Cu, Zn) and a single element standard for Se (VHG LSEN-100). Data were acquired in triplicate and averaged. A coefficient of variance (CoV) was determined from frequent measurements of a sample containing ∼10 ppb of Mn, Fe, Cu, and Zn and ∼1 ppb of Se. An internal standard (Sc, Ge, Bi) continuously introduced with the sample was used to correct for detector fluctuations and to monitor plasma stability. The accuracy of the calibration curve was assessed by measuring NIST reference material (water, SRM 1643f) and found to be within 96% - 105% for all elements.

### Glutathione measurements

The day before the experiment, 2.5 x 10^5^ control, LRP8KO1, or LRP8KO2 HCC1143 cells per well were seeded in duplicate into a 6-well plate (Corning, #07-200-83). The following day, cells were harvested by scraping (2 wells/genotype were combined) and prepared for measurement of total intracellular glutathione (GSH+GSSG) using a glutathione assay kit based on Ellman’s reagent (DTNB, 5,5′-dithio-bis-2-nitrobenzoic acid) according to the manufacturer’s protocol (Cayman Chemical, t#703002). Data are presented as DNTB fluorescence normalized to total protein. Three independent biological replicates were analyzed for each genotype.

### RNA-seq and RT-qPCR

Control and two LRP8KO HCC1143 cell lines were incubated in regular medium or medium containing 200nM Se for 48 hr. Total RNA was extracted using the Monarch Total RNA Miniprep Kit (New England Biolabs, NEB, #T2010S). RNA-seq libraries were constructed using the Swift Rapid Library Prep Kit (Swift Bio #R2096) with NEBNext Poly(A) mRNA Magnetic Isolation Module (NEB #E7490L) according to the manufacturer’s instructions. Samples were sequenced on an Illumina NovaSeq 6000 with PE150 reads. Differential gene expression was analyzed using DESeq2^42^.

The same sets of RNA samples were reverse transcribed using Protoscript II First Strand cDNA Synthesis Kit (NEB #E6560S). cDNA was measured using a QuantStudio 7 Flex Real-time PCR machine (Applied Biosystems) with Power SYBR green PCR master mix (Thermo Fisher Scientific, #4368577). Fold change in mRNA levels were determined using the 2-delta cycle threshold method, normalized to actin (ACTB) mRNA. IDT predesigned PrimeTime primer pairs were purchased, and sequences are as follows: GPX4_1, 5’-GCTGTGGAAGTGGATGAAGA-3’; GPX4_2, 5’-CAGCCGTTCTTGTCGATGA-3’; ACTB_1, 5’-ACAGAGCCTCGCCTTTG-3’; ACTB_2, 5’-CCTTGCACATGCCGGAG-3’.

### Ribosome profiling

Cells were washed once in ice-cold PBS and lysed without cycloheximide in ribosome buffer (20 mM Tris•Cl pH 7.4, 150 mM NaCl, 5 mM MgCl2, 1 mM DTT) with 1% v/v Triton X-100 and 25 U / ml Turbo DNase I [McGlincy et al 2017]. RNA content of cell lysates was quantified using Quant-iT RiboGreen RNA assay kit. Ribosome footprints were generated using Nuclease P1 [NEB M0660S], following a protocol to be described elsewhere (Ferguson L et al., unpublished).

300 µL of lysate was adjusted to pH 6.5 with the addition of 21 µL of 300 mM Bis-Tris. Ribosome footprinting was carried out by adding 900 U Nuclease P1 per 30 µg of total RNA, and nuclease treatment proceeded for 1 hr at 37 °C as described in Ferguson L et al. (unpublished). Nuclease-treated lysates were overlayed on 1M sucrose cushions and ribosomes were sedimented by centrifugation in a TLA-110 rotor at 100,000 RPM for 1 hr. RNA was harvested from the ribosomal pellet using a DirectZol RNA Purification Kit according to the manufacturer’s instructions. Footprints were separated by denaturing electrophoresis in a 15% polyacrylamide gel containing 7 M and 0.6 X TBE. Gel slices containing RNA between 30 and 40 nt were excised and the RNA was eluted overnight at room temperature in elution buffer (300 mM NaCl, 10 mM Tris pH 8, 1 mM EDTA, 0.25% v/v SDS) and precipitated by addition of 3 volumes of ethanol followed by 3 hr -80 °C incubation. RPFs were pelleted by a 30 min 20,000 x g centrifugation at 4 °C. The RPF pellet was washed with 70% ethanol once before being resuspended in 10 mM Tris pH 8 and quantifed by NanoDrop.

Sequencing libraries were constructed using the Ordered Two-Template Relay approach for small RNA library construction^43^ in a protocol to be described elsewhere (Ferguson et al., unpublished). 30 to 40 ng of RPFs were used for Ordered Two-Template Relay ribosome profiling (OTTR-RP). The cDNA product from OTTR-RP was quantified by qPCR [McGlincy et al 2017], and deep sequencing libraries were prepared by amplification of cDNA with unique i5 and i7 TruSeq primers in 12 cycles of PCR. Libraries were sequenced on a NovaSeq S4 with 150 base paired-end sequencing, and only the R1 reads were used for analysis. Adapters were trimmed using Cutadapt v3.2^44^. Human rRNA reads were removed by mapping to several human rRNA sequences and loci (U13369.1, NR_145819.1, NR_146144.1, NR_146151.1, NR_146117.1, X12811.1, ENST00000389680.2, ENST00000387347.2, NR_003287.4, NR_023379.1, NR_003285.3, NR_003286.4) by Bowtie v1.0.0^45^ allowing for up to three mismatches. Human tRNA reads were removed by tRAX v1.0^46^. Reads that did not align to rRNA or tRNA references were then aligned with Bowtie by mapping to the GENCODE v37 protein coding transcripts, and both gene-level quantification and multi-mapping read handling were performed by RSEM v1.1^47^. Up to two mismatches were allowed while mapping RPFs to protein coding transcripts, and protein coding gene quantification was restricted to those reads which aligned to the 15^th^ to the -10^th^ codon of the coding sequence, and any read with more than 250 alignments were ignored.

### Pause detection and translation efficiency measurements

Pause detection and coverage was performed on reads which aligned to the NCBI RefSeq isoform for each selenoprotein transcript. The 5′ positions of alignments were used to determine the start of the A-site codon by adding 15 to frame-0 alignments, 16 to frame-2 alignments, or 17 to frame-1 alignments. Collapsed codon counts for a gene were rescaled by dividing each codon by the average codon counts, and the replicates were collapsed by averaging the rescaled codon density for each codon. For each selenoprotein, the upstream density was taken by summing the densities from the start codon to the SEC UGA codon, and the downstream density was taken by summing the densities from the codon after the SEC UGA to the stop codon. Upstream and downstream densities were rescaled by the length of the respective regions. Translation efficiency estimates were made by DESeq2 following the deltaTE approach using ribosome profiling and aforementioned RNA-seq gene-level counts made by RSEM^48^. RNA-seq reads were requantified using the same procedure for ribosome profiling prior to translation efficiency estimates.

### Spheroid culture

2500 cells in 100 µL RPMI media supplemented with complete serum were seeded in 96-well Black/Clear Round Bottom Ultra-Low Attachment Spheroid Microplates (Corning, #4515). Cells were incubated at 37 °C for 30 min, followed by the addition of 100 µL RPMI media containing 2% Matrigel (Corning, #354234). Plates were centrifuged at 750g for 15 min and grown at 37 °C for 2 days. For IKE treatment, 100 µL RPMI media was slowly removed without disturbing the spheroid. Another 100 µL RPMI media containing 1% Matrigel, 18 µM IKE, and 60 nM SYTOX Green dye was added back into each well. Spheroids were imaged in the IncuCyte S3 with a 10x objective.

### BONCAT for analysis of GPX4 translation

Cells were grown in 15-cm dishes until 70% confluent. RPMI media was replaced with methionine-free RPMI (Gibco, #A1451701) and cells were cultured for 1 hr to remove existing methionine. 25 μM Click-I AHA (L-Azidohomoalanine, Thermo Fisher, #C10102) was added to the media, cells were incubated at 37 °C for 2 hr, and cells subsequently washed 3x with cold PBS. Cells were lysed with RIPA buffer and sonication. Protein was quantified using the BSA assay, equal protein amounts were collected across the samples and subjected to 25 μM DBCO-PEG4-biotin (Sigma-Aldrich, #760749) at room temperature for 30 min. Acetone precipitation was performed to remove any excess DBCO-biotin in the sample. The protein pellet was then resuspended in 2% SDS and diluted with RIPA buffer. Meanwhile, 25 μL per sample of NeutrAvidin Agarose resins (Pierce, # 29201) were washed 3x in RIPA buffer and distributed to the lysates. Resin and lysates were incubated at 4 °C overnight. The beads were washed 4x with RIPA containing 400mM NaCl and were boiled in RIPA (with 1x Laemmli buffer and 25mM Biotin) for 5 min at 95 °C. One third of the lysates were separated on 4–20% polyacrylamide gradient gels for western blot analysis. The remaining two thirds of cell lysates were separated on another gradient gel, and 10-25 kDa regions were cut, extracted, and processed for MS analysis. Briefly, gel bands were destained with 10 gel volumes of 50 mM ammonium bicarbonate (ABC) pH 8.0 and 50% acetonitrile (ACN) with agitation for 45 min. The solution was removed and replaced with 100% ACN to ensure complete gel dehydration.

Modified sequencing grade trypsin (Promega #PRV5111) was diluted to 50 ng/µL using 50 mM ABC and added to gel pieces at approximately equivalent gel volumes and incubated on ice for 1 hr. Another 1x gel volume of trypsin was then added to samples and digested overnight at 37 °C. Digests were quenched with 50% acetonitrile, 5% trifluoroacetic acid, and vortexed. Digested peptides were transferred into fresh Eppendorf tubes, two additional extraction steps were performed, and peptides were combined and then dried to completion in a speed vac. Peptides were resuspended in 1% formic acid and separated on an Easy nLC 1000 UHPLC equipped with a 15 cm nanoLC column. Using a flow rate of 300 nL/min, the linear gradient was 5% to 35% over B for 90 min, 35% to 95% over B for 5 min, and 95% hold over B for 15 min (solvent A: 0.1% formic acid (FA) in water, solvent B: 0.1% FA in ACN). Peptide identities and relative abundances were determined using Proteome Discoverer 2.4. Ion chromatograms were extracted using Xcalibur Qual Browser for each peptide of interest with a mass tolerance of 0.5 Da. Results were normalized against the average control value and are represented as average ± SD of three replicates.

### Statistical analysis and reproducibility

All figures, including Western blots, dose-response curves, and panels are representative of at least three biological replicates unless stated otherwise. *P*-values for pairwise comparisons were calculated using the two-tailed t-test. For comparison across multiple experimental groups, *p*- values were calculated using one-way ANOVA.

## EXTENDED DATA FIGURE LEGENDS

**Extended Data Fig. 1:**
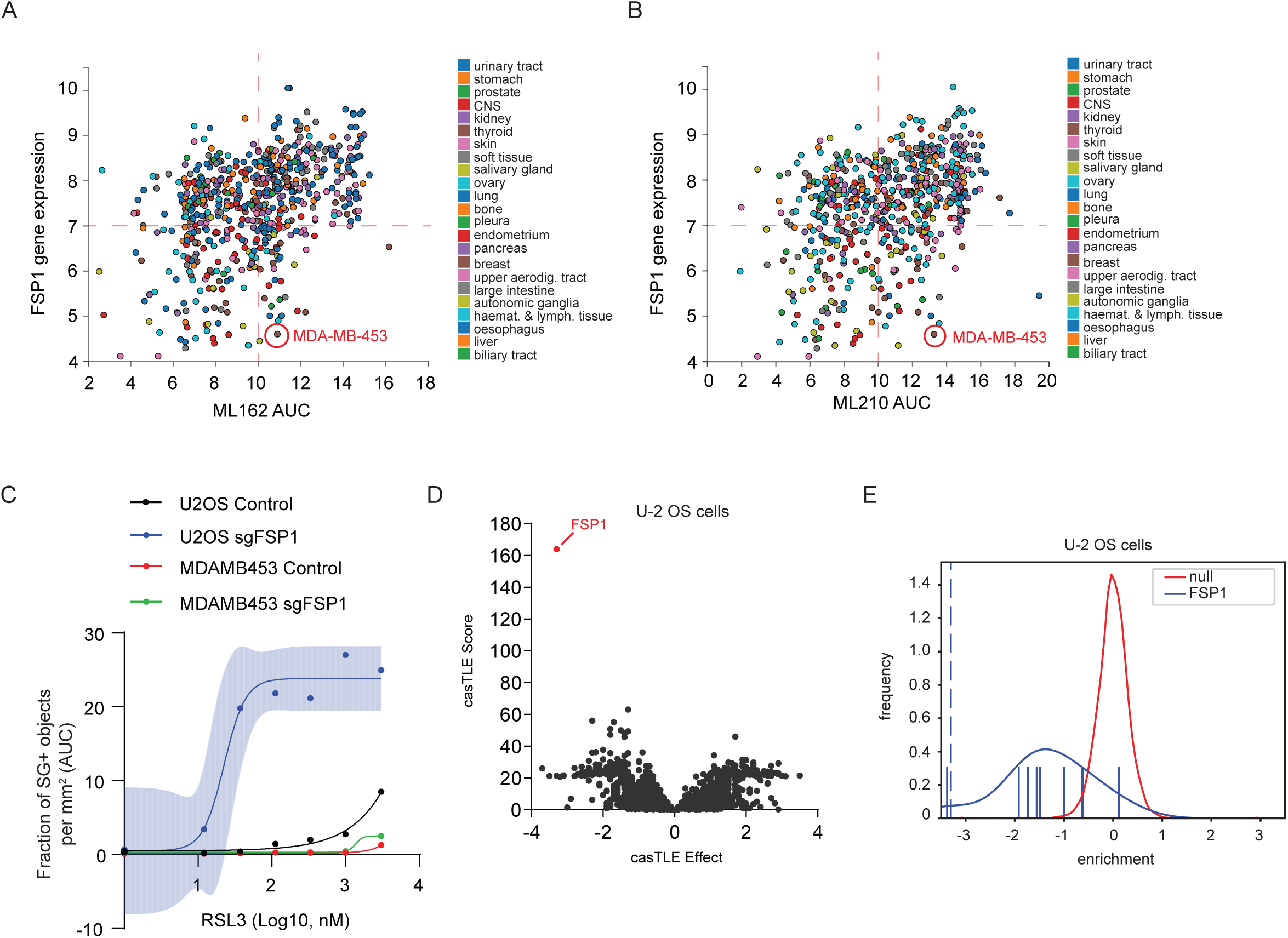
Cell-specific role for FSP1 in ferroptosis. **a-b**, Summary of FSP1 gene expression and ML162 (**a**) and ML210 (**b**) sensitivity in various cancer cells. Data were mined from the CTRP database. **c**, Dose response of RSL3-induced cell death of control and FSP1KO in U-2 OS and MDA-MB-453. Shading indicates 95% confidence intervals for the fitted curve, and each data point is the average of three replicates. **d**, Gene effects and gene scores calculated for individual genes analyzed in the genome-wide CRISPR screen of U-2 OS cells. . **e**, Histogram of FSP1 from casTLE analysis showing the negative enrichment of FSP1 compared to controls.

**Extended Data Fig. 2:**
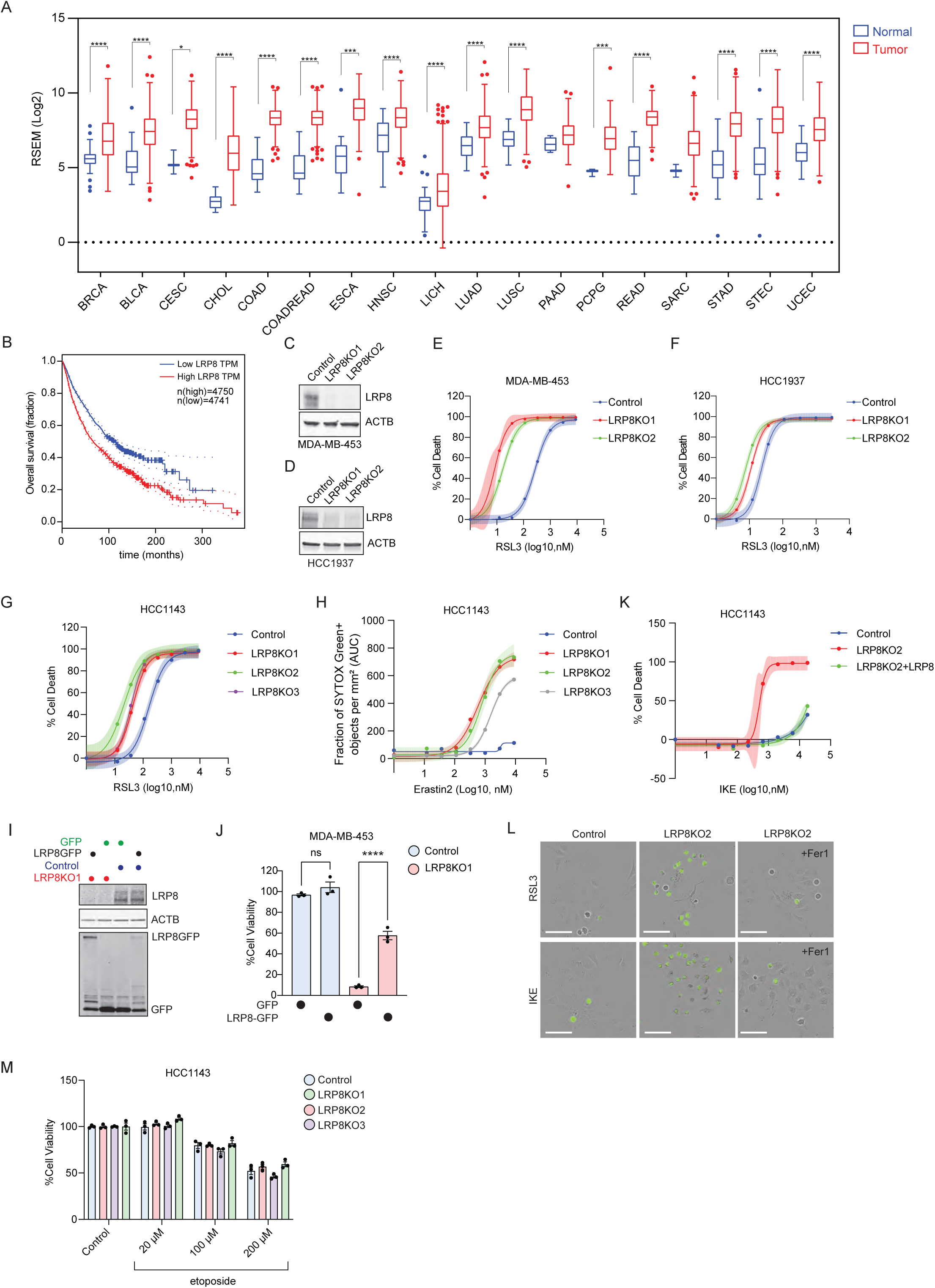
Analysis of LRP8 in cancers and as a ferroptosis resistance factor. **a**, Relative expression of LRP8 in normal tissue and tumors from various primary sites. Plotted data were mined from the TCGA database. LRP8 is highly expressed in tumors across many cancer types. **b**, Kaplan-Meier survival curve showing that patients with higher expression of LRP8 have poorer survival rate. Plotted data were mined from the TCGA database. **c**, Western blot to validate LRP8KO in MDA-MB-453 cells. Two single-clonal LRP8KO lines generated from different sgRNAs were selected and compared with parental cells stably expressing Cas9. **d**, Western blot to validate LRP8KO in HCC1937 cells. Two single-clonal LRP8KO lines generated from the same sgRNA were selected and compared with control cells expressing sgSAFE. **e**, Dose response of RSL3-induced cell death of control and LRP8KO MDA-MB-453 cells. **f**, Dose response of RSL3-induced cell death of HCC1937 control and LRP8KO cells. **g**, Dose response of RSL3-induced cell death of control and LRP8KO HCC1143 cells by cell-titer glow assay. **h**, Dose response of Erastin2-induced cell death of control and LRP8KO HCC1143 cells. **i**, Validation of LRP8 overexpression in LRP8KO MDA-MB-453 cells. **j**, Cell viability of MDA-MB-453 control and LRP8KO cells stably expressing GFP or LRP8-GFP subjected to 100 nM RSL3 for 24 hr. **k**, Dose response of IKE-induced cell death of control, LRP8KO, and LRP8KO expressing LRP8-GFP. **l**, Live-cell imaging of control and LRP8KO cells incubated with SYTOX Green and treated with 100 nM RSL3 alone or co-treated with 2 µM Fer1. Scale Bar, 100 µm. **m**, Cell viability of HCC1143 control and LRP8KO cells upon the treatment of different doses of etoposide. In **e,f**,**g,h,** and **k** shading indicates 95% confidence intervals for the fitted curve and each data point is the average of three biological replicates. Data in **j** and **m** represent mean ± S.E.M. of three biological replicates by two-tailed, unpaired t-test.

**Extended Data Fig. 3:**
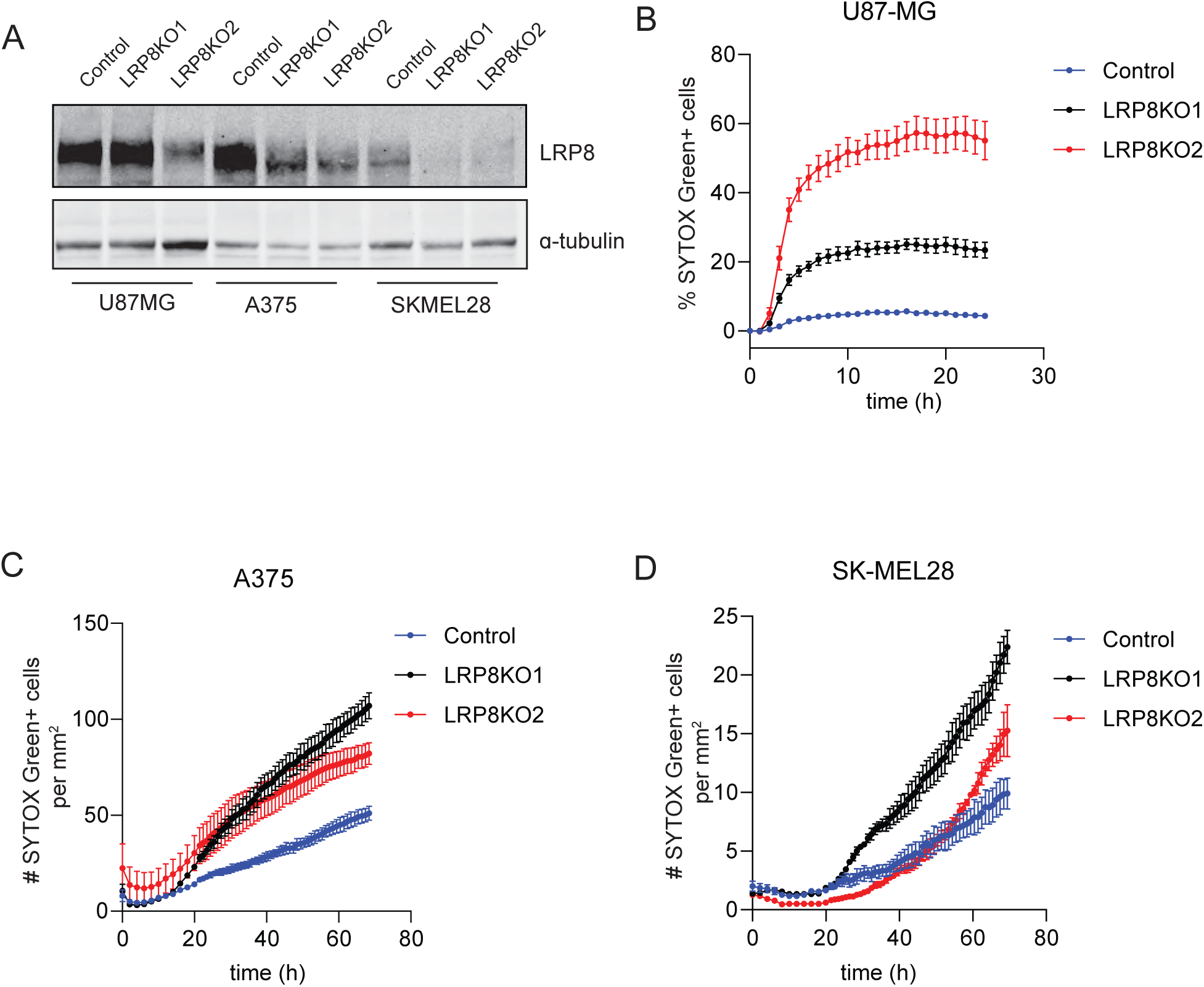
LRP8 promotes ferroptosis resistance in multiple cancer cell lines. **a**, Validation of LRP8KO in multiple cancer cell lines. Westen blot of pools of LRP8KO cells generated from two different sgRNAs and control Cas9-expressing cells. **b-d**, Time-lapse cell death analysis of U87-MG (**b**), A375 (**c**), and SK-MEL28 (**d**) cells treated with RSL3 over 24 hr or 72 hr. Data represent mean ± S.E.M. of three replicates.

**Extended Data Fig. 4:**
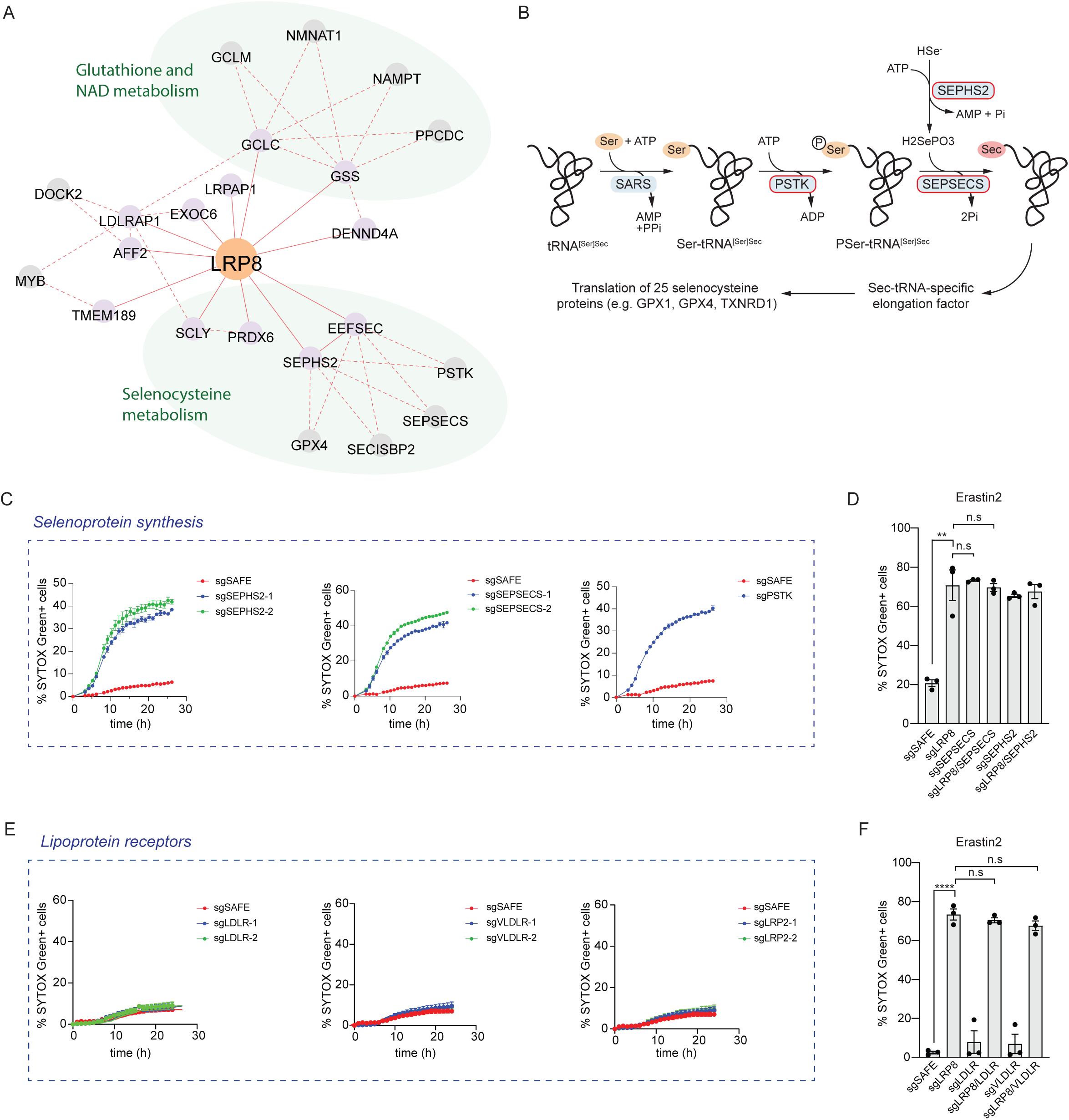
Role of selenium metabolism and lipoprotein receptors in ferroptosis. **a**, Coessentiality network analysis for LRP8 in pan-cancer context by FIREWORKS. Two primary modules, (1) glutathione and NAD metabolism and (2) selenocysteine metabolism, are positively associated with LRP8. **b**, Schematic of selenocysteine synthesis. **c**, Time-lapse cell death analysis of control and CRISPR KO of genes from the selenoprotein synthesis pathway [SEPHS2 (left), SPESECS (mid), PSTK (right)]. Cells were treated with 100 nM RSL3 for 24 hr. **d,** Quantification of the percentage of SYTOX green positive cells (dead cells) of control and single or double KO of indicated genes related to selenocysteine metabolism. **e,** Time-lapse cell death analysis of control and CRISPR knockout of genes from lipoprotein receptor superfamily [LDLR (left), VLDLR (mid), LRP2 (right)]. Cells were treated with 100 nM RSL3 for 24 hr. **f,** Quantification of the percentage of SYTOX green positive cells (dead cells) of control and single or double KO of indicated genes related to lipoprotein receptor superfamily. Data in **d, f** represent mean ± S.E.M. of three biological replicates by two-tailed, unpaired t-test.

**Extended Data Fig. 5:**
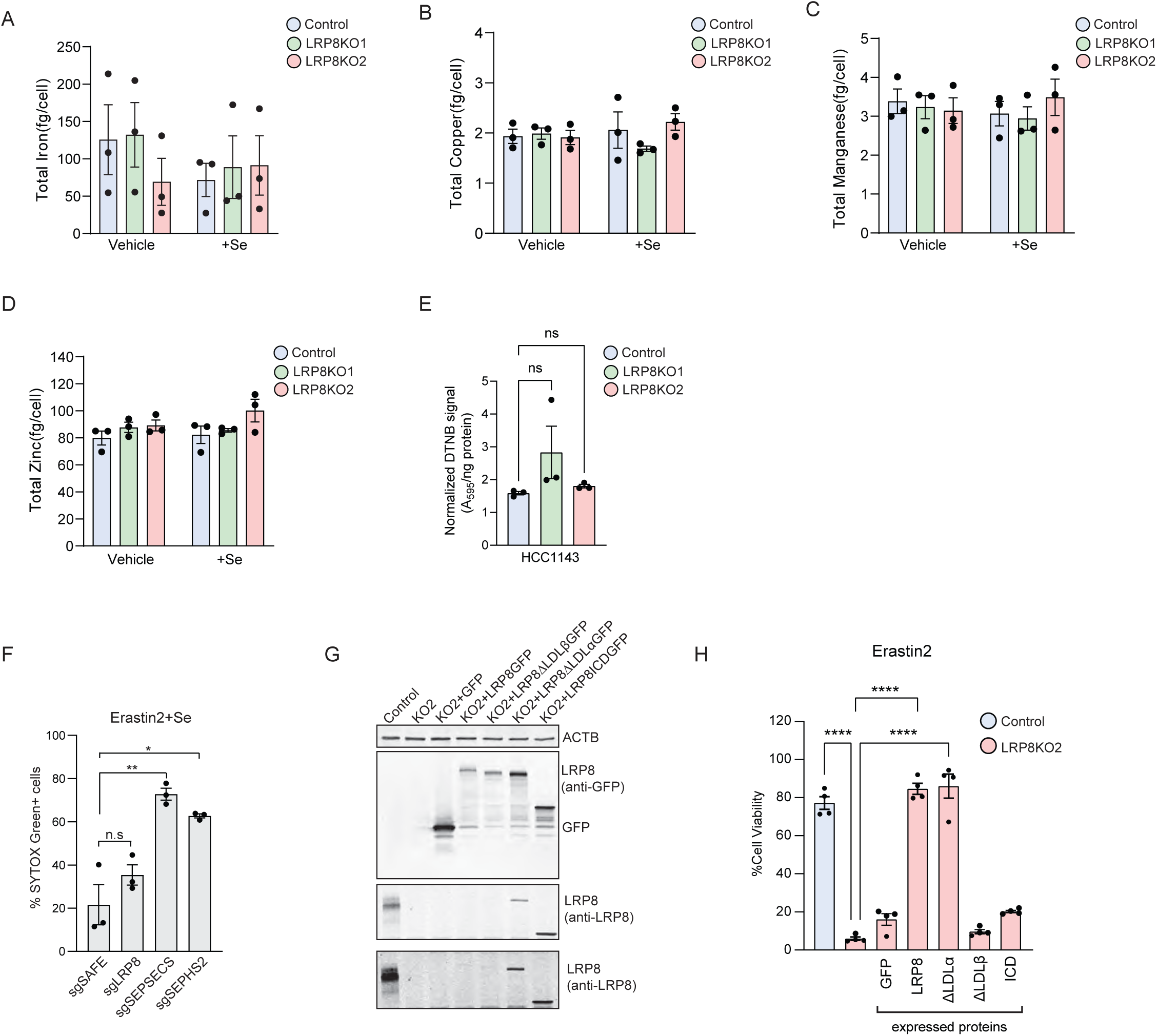
Measurement of metals in LRP8 knockout cells. **a-d**, Total levels of iron (**a**), copper (**b**), manganese (**c**), and zinc (**d**) in control and LRP8KO cells alone or the presence of 200 nM Se by ICP-MS. **e**, Measure of the glutathione level (DNTB) in HCC1143 control and LRP8KO cells. **f**, Quantification of SYTOX green positive cells in KO cells of indicated genes subjected to 3 µM Erastin2. **g**) Immunoblot showing the expression of GFP tagged LRP8 wildtype or truncation mutants in LRP8KO HCC1143 cells. **h**, Cell viability of control cells or LRP8KO cells stably expressing indicated LRP8 mutants in the presence of 3 µM Erastin2 for 24 hr. All data represent mean ± S.E.M. of three biological replicates by one-way ANOVA (**a-d,h**) or paired t-test (**e,f**).

**Extended Data Fig. 6:**
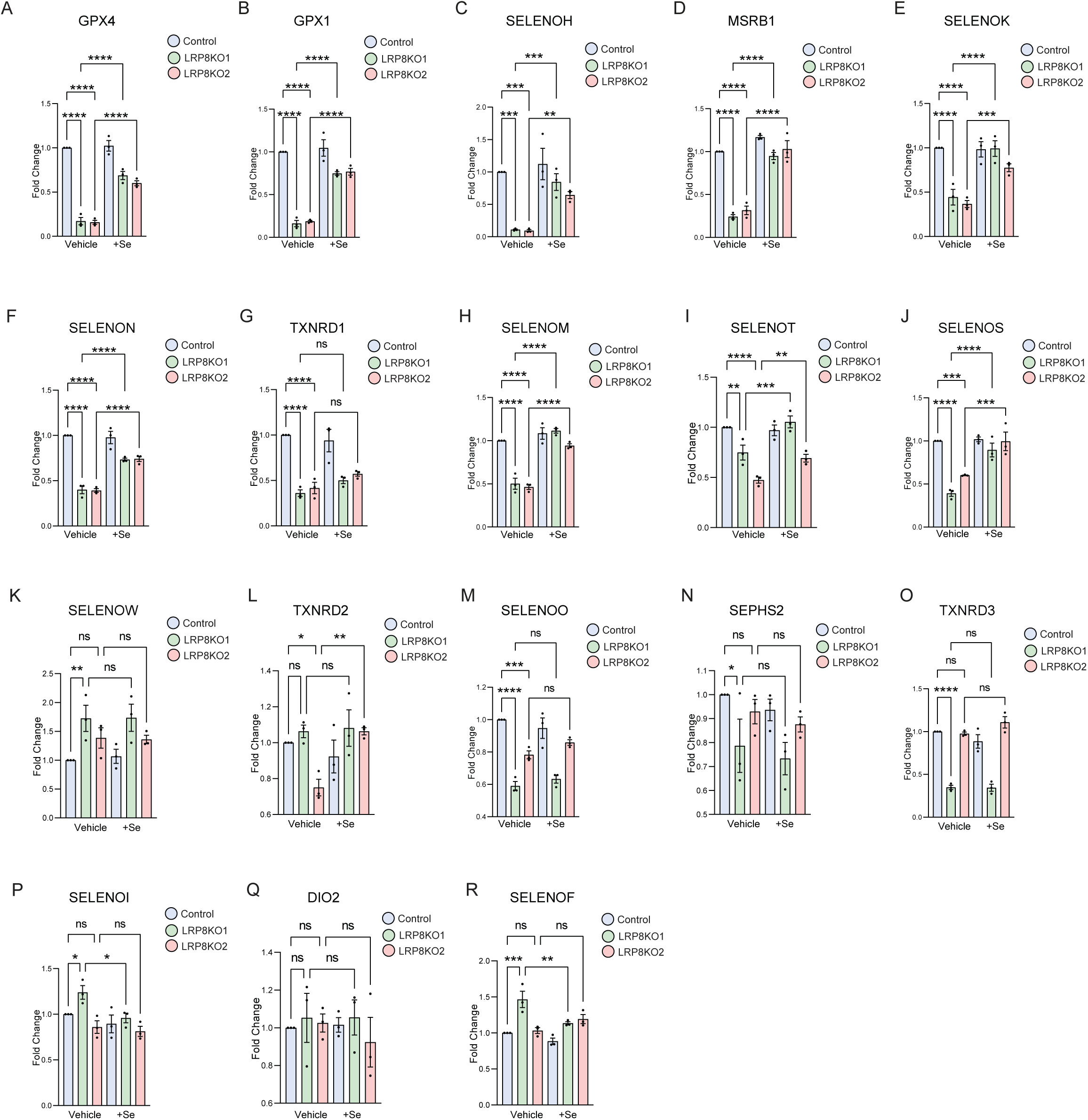
LRP8 and selenium affect selenocysteine protein levels. **a-r**, Quantification of changes in the selenocysteine protein levels of control and LRP8KO cells with or without 200 nM Se supplement. Fold-change in individual selenocysteine proteins were plotted in a separated bar graph. Data represent mean ± S.E.M. of three biological replicates by one-way ANOVA.

**Extended Data Fig. 7:**
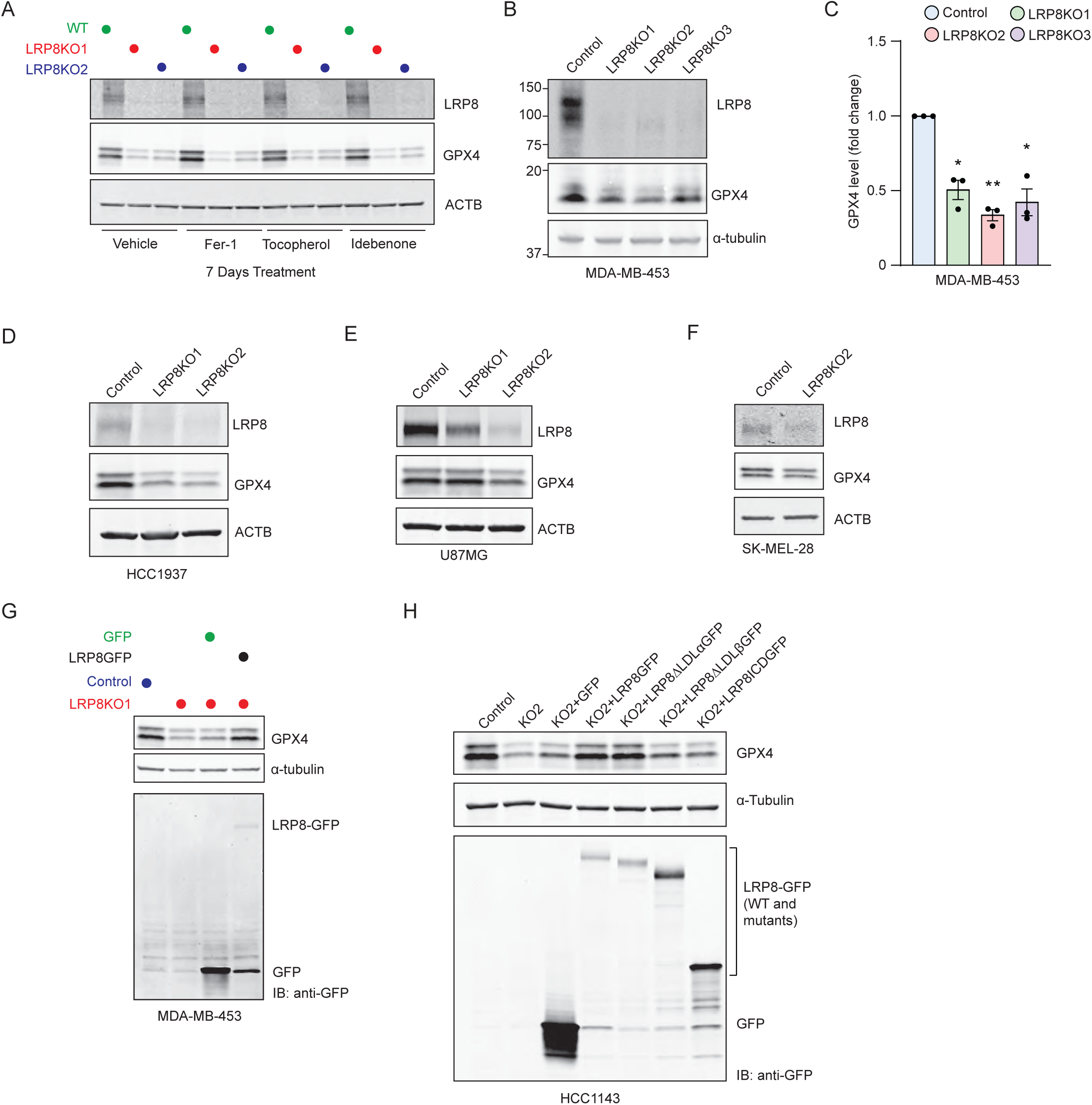
Loss of LRP8 results in reduced GPX4 levels in multiple cancer cell lines. **a**. Representative western blot of HCC1143 cell lysates subjected to various antioxidants for seven days. **b,** Representative western blot of GPX4 in MDA-MB-453 cell lysates from multiple clonal LRP8KO lines. **c**, Quantification of GPX4 levels from **b** (three independent replicates). **d-f**, Representative western blot of GPX4 in HCC1937 (**d**), U87-MG (**e**), SKMEL38 (**f**) LRP8KO cells. **g,h**, Western blot of GPX4 in MDA-MB-453 (**g**) and HCC1143 (**h**) LRP8KO cells stably expressing LRP8 wildtype or various truncation mutants. Data in **b** represent mean ± S.E.M. of three biological replicates by paired t-test.

**Extended Data Fig. 8:**
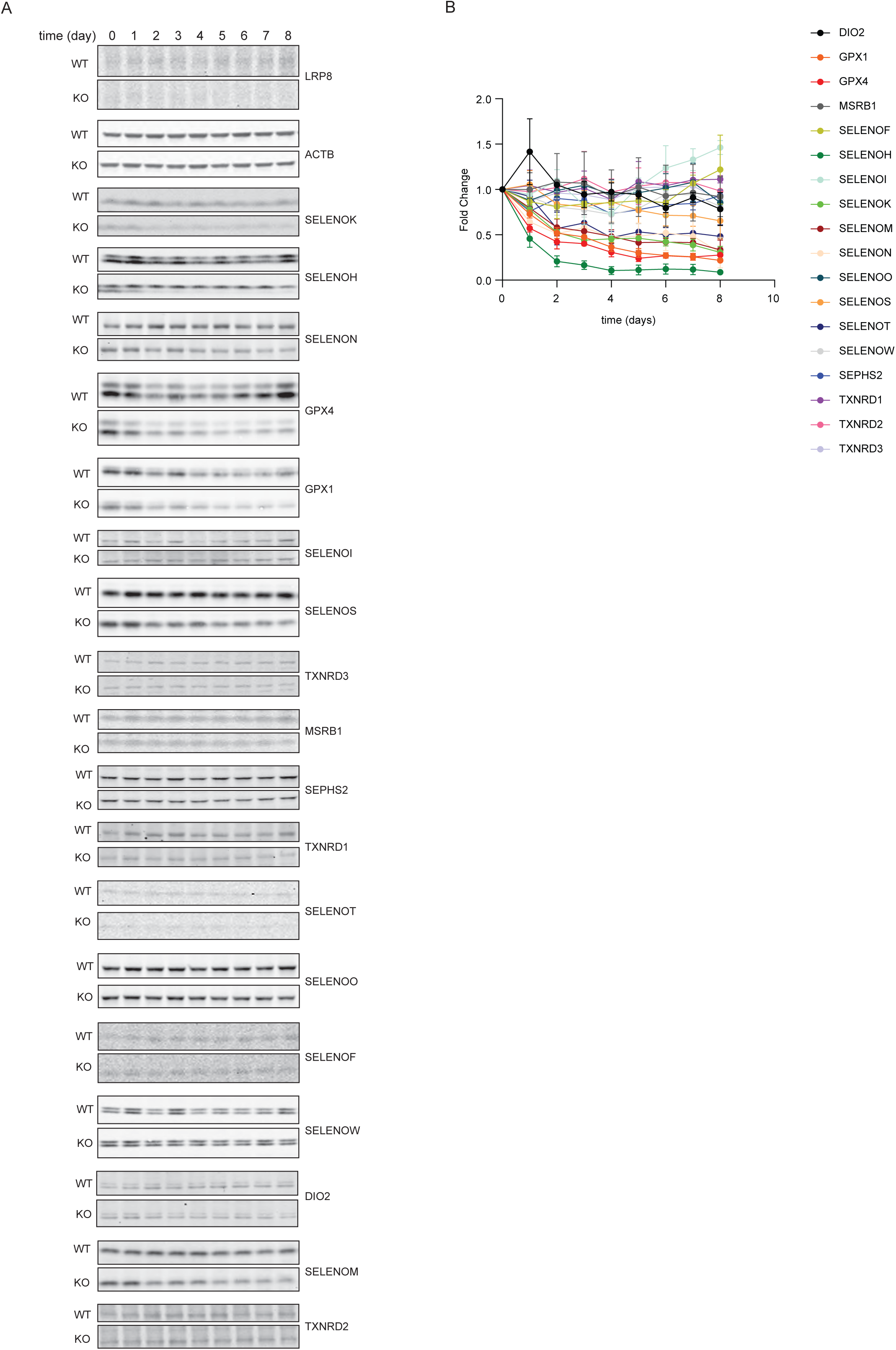
Regulation of selenocysteine proteins. **a**, Representative western blots showing changes in selenocysteine protein levels upon selenite withdrawal. **b**, Quantification of selenocysteine protein levels from **a** (*n* = 3 independent experiments). The level of each protein was first normalized to that of ACTB from the same lysate sample and subsequently normalized to that of LRP8KO from Day 0.

**Extended Data Fig. 9:**
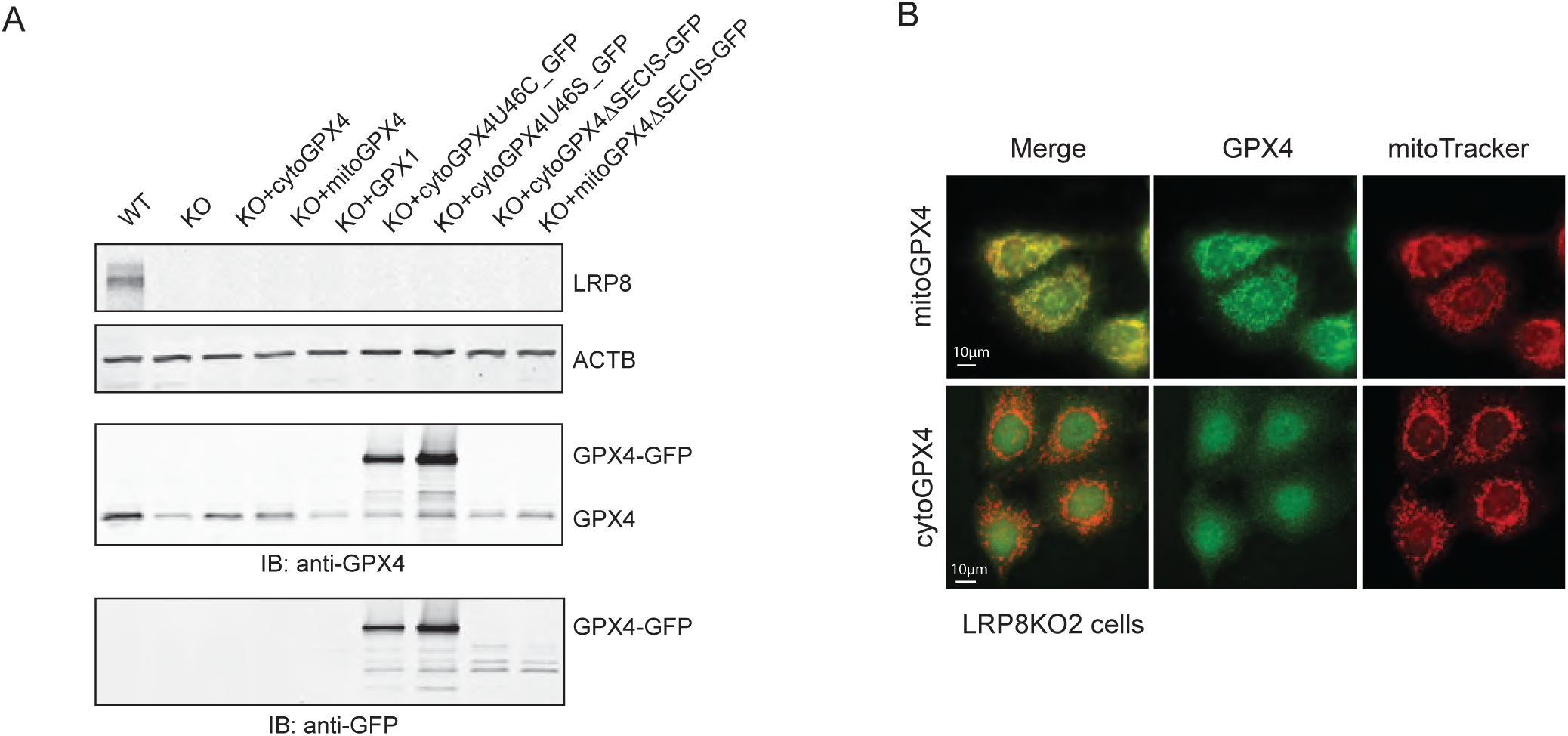
GPX4 overexpression rescues LRP8KO from ferroptosis. **a**, Validation of GPX4 wildtype and mutant expression in HCC1143 LRP8KO cells. **b**, Immunofluorescence of GPX4 (green) in LRP8KO cells stably expressing sGPX4 or lGPX4. Mitochondria were labeled by MitoTracker Deep Red. Scale bars, 10 μm.

**Extended Data Fig. 10:**
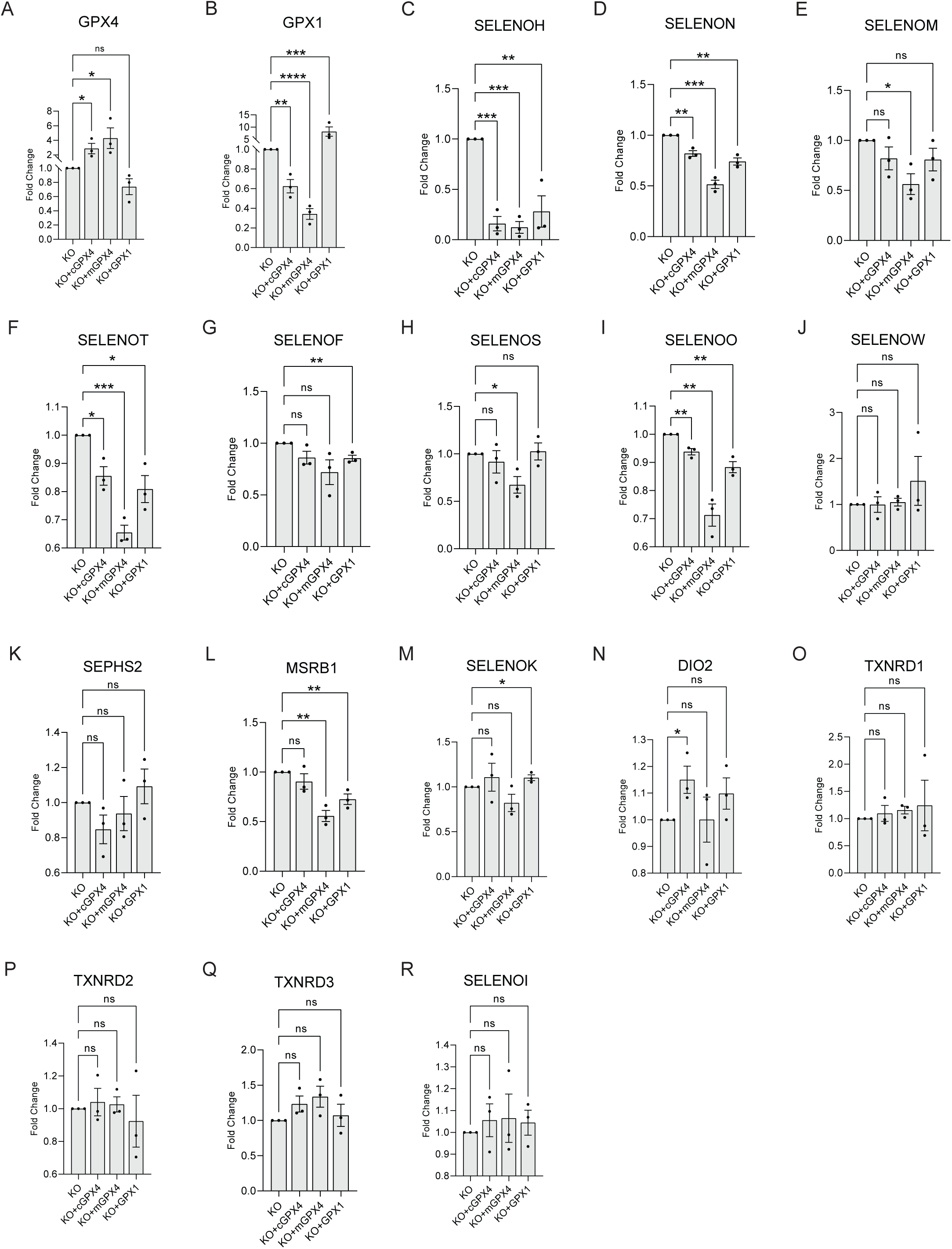
GPX4 overexpression affects on selenocysteine protein levels. **a-r**, Quantification of Western blots ofselenocysteine protein levels in LRP8KO cells stably overexpressing sGPX4, lGPX4, or GPX1. Panels represent mean ± S.E.M. of three biological replicates analyzed by two-tailed unpaired t-test, except for statistical analysis between KO and KO+cGPX4; KO and KO+mGPX4 in (**a**), as well as between KO and KO+GPX1 in (**b**), which used one-tailed, unpaired t-test.

**Extended Data Fig. 11:**
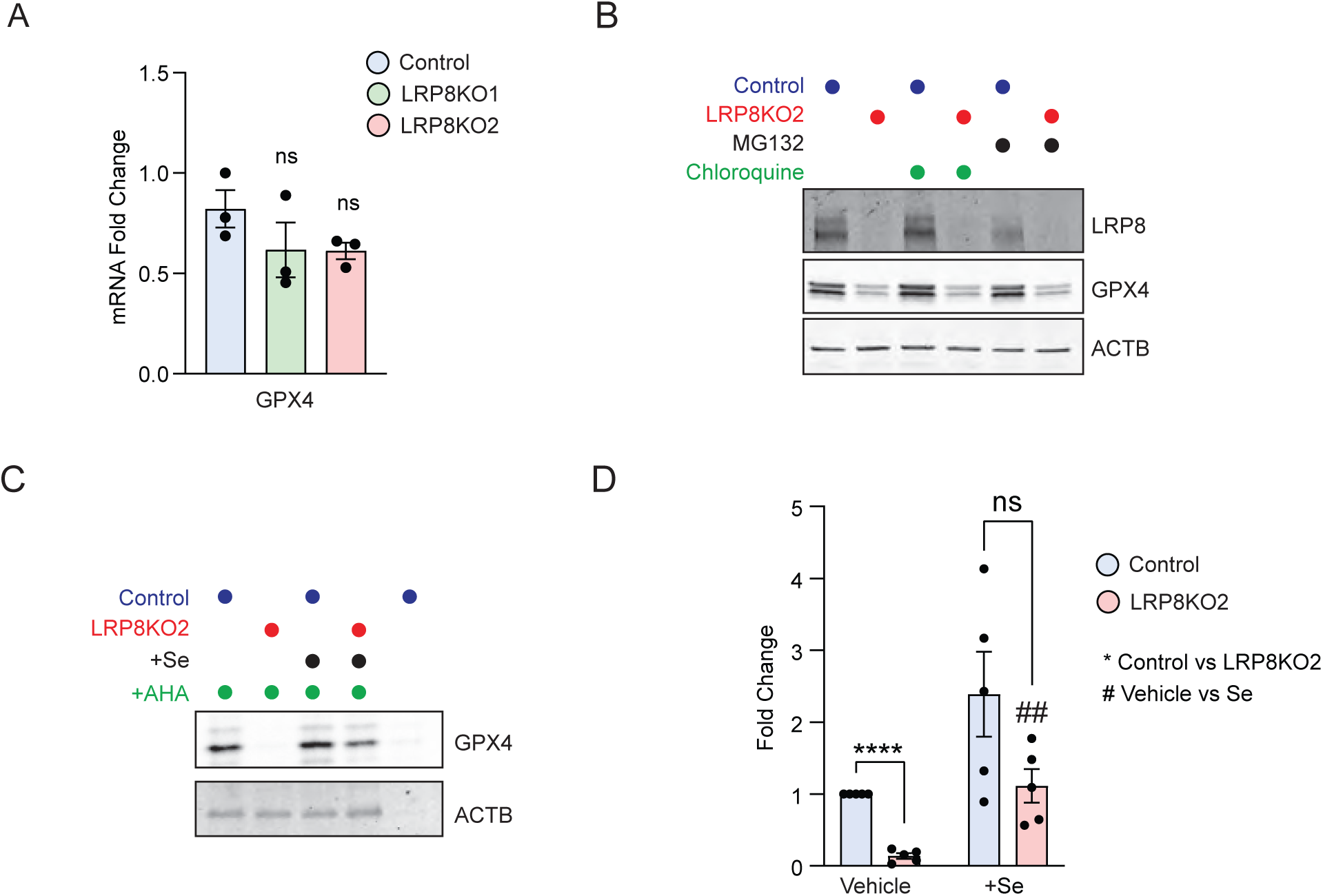
LRP8KO impacts GPX4 translation but not transcription or protein turnover. **a**, Relative GPX4 mRNA levels in control and LRP8KO cells as measured by quantitative PCR. Data represent mean ± S.E.M. of three biological replicates by two-tailed, unpaired t-test. **b**, western blot of GPX4 in control and LRP8KO cells following 10 μM MG132 or 20 μM Chloroquine treatment for 24 hr. **c**, Representative western blots of AHA labeled, newly translated GPX4 in control and LRP8KO cells. **d**, Quantification of GPX4 protein levels from **d**. Data represent mean ± S.E.M of five biological replicates by paired t-test.

**Extended Data Fig. 12:**
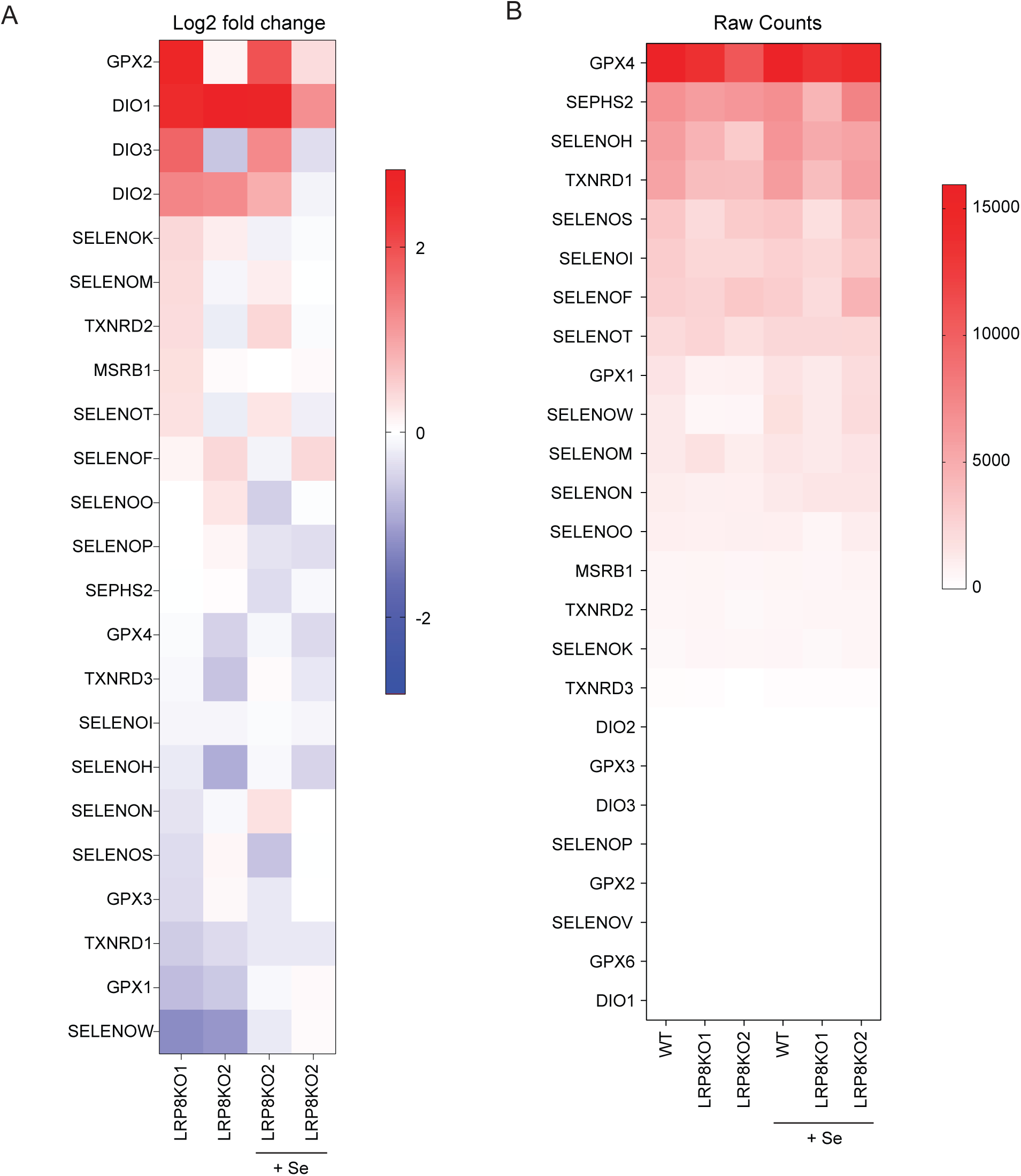
Analysis of gene expression in LRP8KO cells. **a**, Gene expression profiles of 23 human selenocysteine genes detected in control and LRP8KO HCC1143 cells with or without pre-treatment with 200 nM Se for 24 hr. Gene expression in LRP8KO cells is represented as log2 fold change relative to that of control cells from the same condition. **b**, Raw sequence counts of the 23 selenocysteine genes from RNA-seq.

**Extended Data Fig. 13:**
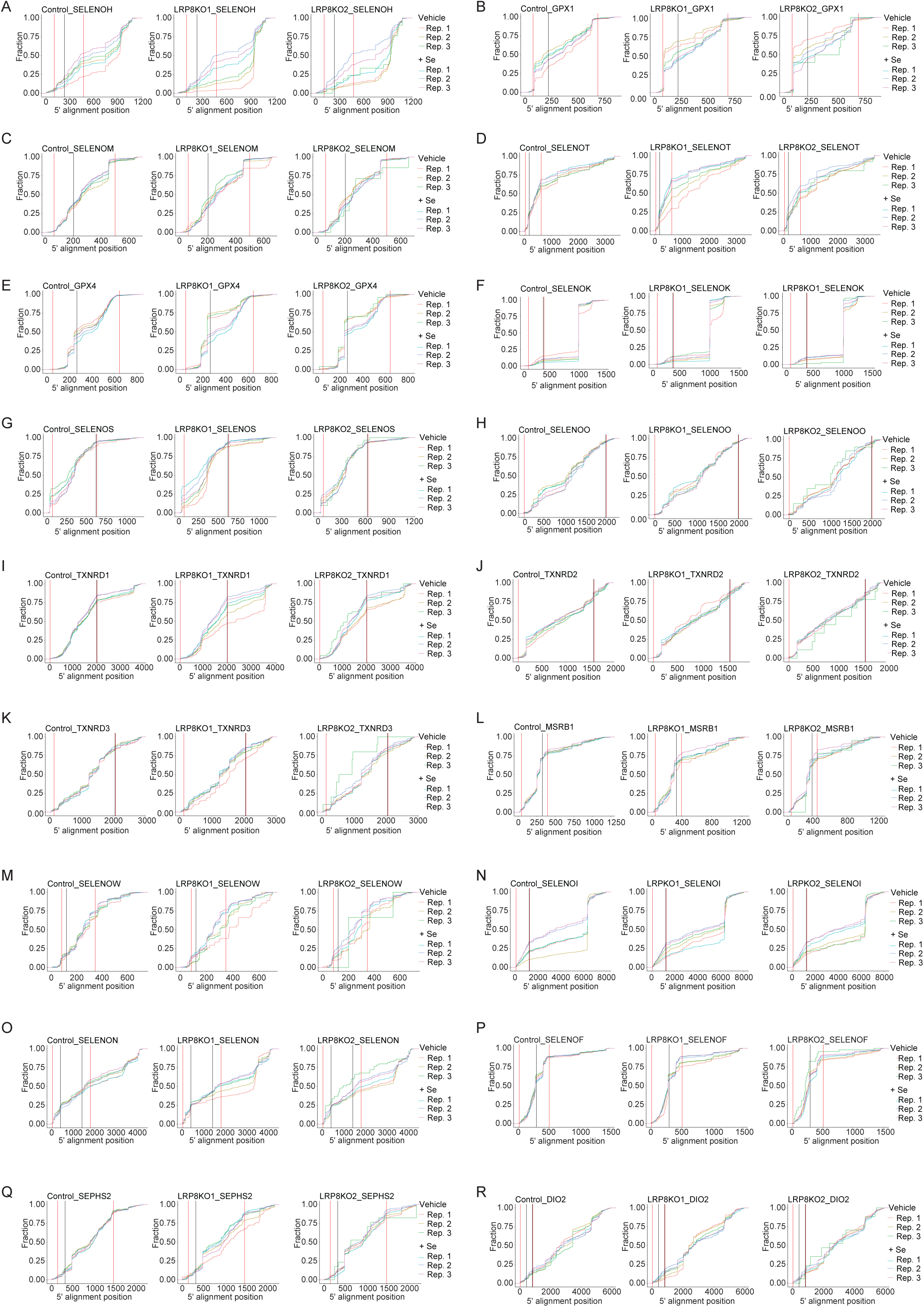
Fraction plots for individual genes encoded selenoproteins. Cumulative frequency of RPFs of 18 SEC transcripts in control and two LRP8KO cell lines incubated with vehicle or Se.

**Extended Data Fig. 14:**
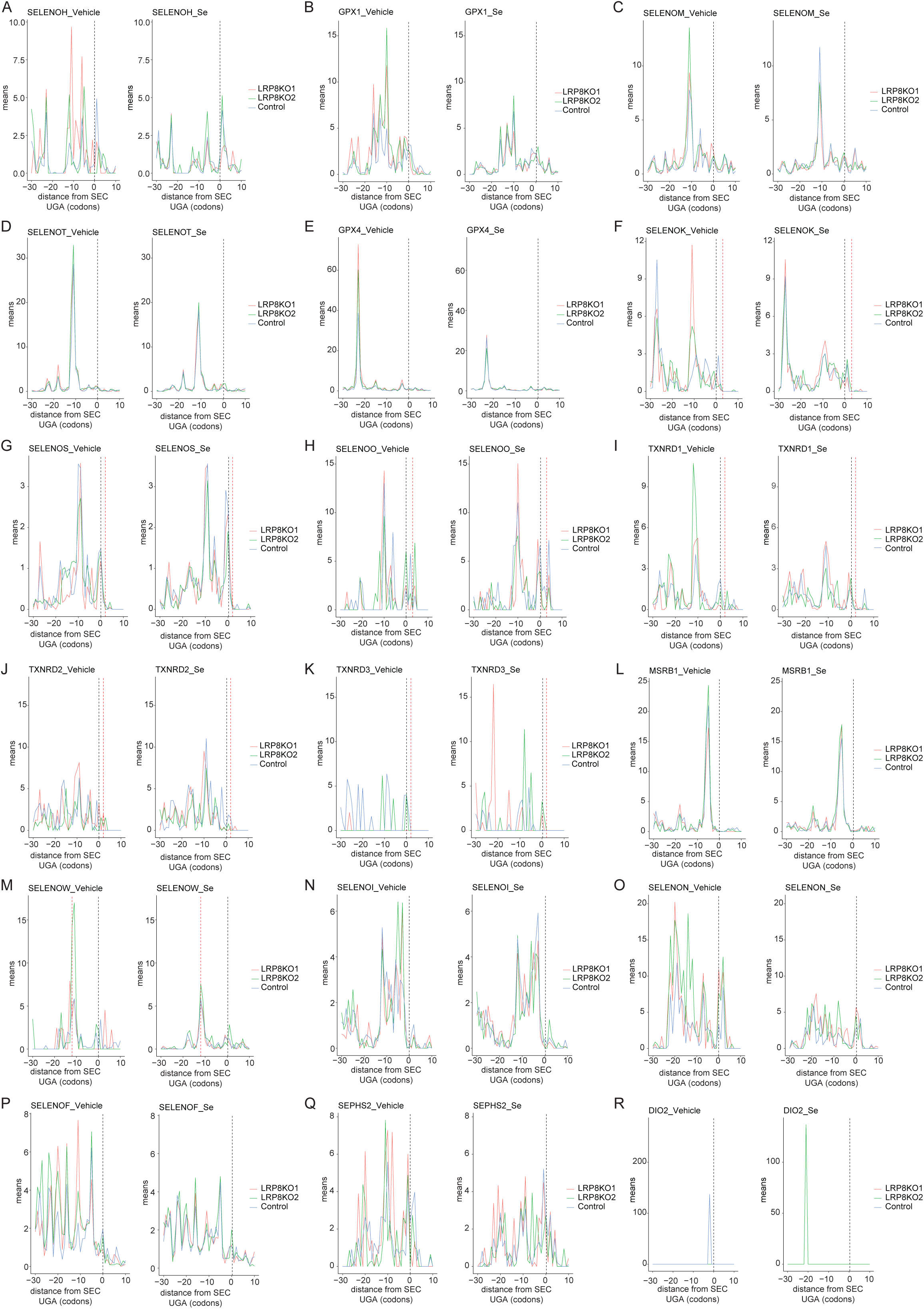
Upstream-ribosome capture for individual genes encoded selenoproteins. Ribosome footprints at the 5’ end of the SEC codon on 18 SEC transcripts from control and two LRP8KO cell lines incubated with vehicle or Se.

## SUPPLEMENTARY TABLES

**Supplementary Table 1: U-2 OS CRISPR-Cas9 synthetic lethal screen data**

Full casTLE results of genome-wide synthetic lethal CRISPR-Cas9 screen in U-2 OS cells.

**Supplementary Table 2: MDA-MB-453 CRISPR-Cas9 synthetic lethal screen data**

Full casTLE results of genome-wide synthetic lethal CRISPR-Cas9 screen in MDA-MB-453 cells.

**Supplementary Table 3: RNA-seq data**

RNA-seq expression data for control and LRP8KO cells incubated with vehicle or Se.

**Supplementary Table 4: Ribosome profiling data**

Translation Efficienct (TE) data for control and LRP8KO cells incubated with vehicle or Se.

## Notes

### Competing Interest Statement

S.J.D. is a co-founder of Prothegen Inc. and a member of the advisory board for Ferro Therapeutics and Hillstream Biopharma. S.J.D. and J.A.O. have patent applications related to ferroptosis.

